# Computational Fluid Particle Dynamics (CFPD)-Based Virtual Next Generation Impactor (vNGI) to Predict the Aerodynamic Particle Size Distribution (APSD) of Respiratory Drug Delivery Products: Toward New Approach Methodologies (NAMs) in Inhaler Performance Evaluation

**DOI:** 10.64898/2026.06.29.735263

**Authors:** Abhijeet Sunilkumar Patil, Yu Feng

## Abstract

The Next Generation Impactor (NGI) is one of the regulatory gold standards for characterizing aerodynamic particle size distributions (APSDs) of orally inhaled drug products (OIDPs); however, its reliance on complex, resource-intensive *in vitro* testing under tightly controlled environmental conditions limits experimental flexibility and introduces variability. In alignment with the growing regulatory emphasis on New Approach Methodologies (NAMs) for drug development, this study presents a rigorously validated computational fluid particle dynamics (CFPD)-based virtual NGI (vNGI) as an *in silico* method complementary to conventional testing. The vNGI replicates a significant portion of the NGI geometry and airflow physics, enabling high-resolution spatiotemporal analysis of aerosol transport and deposition mechanisms that are otherwise inaccessible experimentally. A comprehensive verification and validation framework was implemented, including mesh and particle independence studies, turbulence model assessment, and comparison of stage-wise deposition efficiencies with available *in vitro* data at 30 L/min. The model’s capabilities were further extended to low and high flow rates, and two bio-relevant mouth-throat models and polydisperse particle-laden aerosol were added. The model demonstrates strong predictive capability for a few stages and provides mechanistic insight into discrepancies in other stages, depending on the type of analysis. Importantly, this work establishes the vNGI as a fit-for-purpose according to NAM by (i) defining a clear context of use for APSD prediction and inhaler performance evaluation, (ii) capturing physically and biologically relevant air-particle interactions, and (iii) demonstrating technical robustness and reproducibility through systematic validation. The platform can potentially further enable simulation of environmental and physiological conditions, such as humidity effects, that are difficult to control experimentally, thereby improving human relevance and reducing reliance on costly and time-consuming *in vitro* testing. This study positions the vNGI as a scalable, regulatory-aligned NAM capable of supporting early-stage drug-device combination product development, device optimization, and an alternative bioequivalence assessment, contributing to ongoing efforts to enhance predictive performance, reduce experimental burden, and transition toward human-centric, inhalation product evaluation.

## 1. Introduction

The inhalation pharmaceutical industry follows the product-specific guidelines (PSG) developed by the regulatory authority to develop and launch products in the regulated market. These guidelines consist of a few *in vitro* and *in vivo* studies, which mainly compare the qualitative and quantitative performance of the product [1, 2]. The regulatory decisions for inhalation drug products are based on a large pool of data generated, such as Single Actuation Content (SAC), emitted Aerodynamic Particle Size Distribution (APSD) from the inhaler mouthpiece, spray pattern, plume topography, priming and repriming, spray duration, and velocity, from *in vitro* bioequivalence (BE) studies and a few *in vivo* studies. The pharmaceutical industry spends a lot of time and money conducting these required studies [3]. This could lead to higher product costs and reduced affordability for users. Despite their widespread use, these experimental approaches are resource-intensive and increasingly challenged by regulatory and scientific efforts to improve efficiency, reproducibility, and human relevance in drug development.

In response to the above-mentioned challenges, regulatory agencies such as the U.S. Food and Drug Administration (FDA) have increasingly emphasized the adoption of New Approach Methodologies (NAMs) [4] to support drug development and regulatory decision-making. NAMs encompass a broad range of non-animal approaches, including *in silico* modeling, to improve predictive performance, reduce reliance on traditional testing methods, and enhance human biological relevance [5]. Recent FDA guidance [4] highlights the importance of developing NAMs that are fit-for-purpose, appropriately validated, and capable of addressing clearly defined contexts of use in drug development.

Within the field of orally inhaled drug products (OIDPs), the application of NAMs remains relatively limited, particularly for predicting aerodynamic particle size distribution (APSD), an important determinant of lung deposition and therapeutic efficacy. Specifically, emitted APSD is one of the key parameters that are usually measured via *in vitro* studies, in which the orally inhaled drug products (OIDPs) are tested for understanding the behavior of the drug-device combination product under the test conditions, such as flow rate, and ambient conditions, using different equipment [6]. The next generation impactor (NGI), designated as the USP <601> Apparatus 6 [7] and Apparatus E as per the European Pharmacopeia [8], is used to measure the APSD. The Next Generation Impactor Consortium developed NGI in the mid-90s [9] and the equipment was filed for patent in 2002 [10]. NGI addresses many of the inadequacies of previously developed pharmaceutical impactors, including a wide range of flow rates and applicability to products such as pressurized metered-dose inhalers (pMDIs), dry powder inhalers (DPIs), nasal products, nebulizers, and soft mist inhalers (SMIs).

SMIs such as Respimat® are built to generate a spray with low momentum and fine droplets [11, 12]. The formulation of SMI is aqueous, which means the spray droplet sizes are sensitive to humidity and temperature. Therefore, the product-specific guidelines of SMIs require the emitted APSD measurement to be conducted at a low flow rate of 28.3 L/min or 30 L/min with either high humidity or low temperature [6]. Schuschnig et al. [13] have proposed keeping the NGI below 5 °C for approximately 90 to 120 min, which could be sufficient to conduct the *in vitro* study as per the PSG. However, maintaining such non-ideal ambient conditions is challenging and can introduce additional errors during *in vitro* studies. Dennis et al. [14] observed drug buildup and corrosion under extremely high-humidity conditions. Suppose the product development is for a powdered form of a drug, as in DPI, the recommended flow-rate conditions change. For example, for an active ingredient such as tiotropium bromide, the steady flow rates recommended for DPI [15] are 20 L/min, 39 L/min, and 60 L/min, whereas for a spray inhaler [6], the recommended spray flow rate is 28.3 L/min.

Based on the challenges of the *in vitro* studies mentioned above, research reports suggest and advocate using a computational model to complement experimental studies [16, 17]. The ongoing support activities by regulatory agencies for the use of *in silico* methods, including empirical models and computational fluid dynamics (CFD) models, to minimize the burden on *in vitro* methods are documented in existing publications [16-20]. CFD, and more specifically computational fluid particle dynamics (CFPD) (e.g., CFD-discrete phase model (DPM)), represents a promising *in silico* NAMS tool capable of resolving complex airflow-particle interactions with high spatial and temporal resolution. Such approaches enable detailed investigation of aerosol transport phenomena under controlled and variable conditions, offering advantages in flexibility, cost efficiency, and mechanistic interpretability compared to traditional experimental methods. A few *in silico* studies have been conducted on the Anderson Cascade Impactor (ACI) using CFD [21-25]. For example, Vinchurkar et al. [21] studied the impact of electrostatic charge on the APSD of aerosols across separate stages using simplified inter-stage boundary-condition coupling. Gulak et al. [23] developed and employed a single-jet model for the 8-stage ACI. Instead of simulating every nozzle at every stage, they solved the 2D axisymmetric Navier-Stokes (N-S) equations to predict the flow field for a representative jet, then tracked particles to reconstruct collection efficiency (*CE*) curves and cut-off diameters (*d*_50_) across different flow rates. Flynn et al. [22] developed a 3D CFD-DPM model for ACI and found that high flow rates intensify recirculation vortices, leading to increased wall losses. Dechraksa et al. [25] investigated how the pre-separator influences airflow and particle transport, and maximizes deposition of desirable particles onto the collection plates in ACI. Recently, Mitani et al. [24] moved beyond spherical particles and studied non-spherical particle deposition in a cascade impactor using A validated CFD-DPM with an optimized drag-force model for non-spherical particles.

Despite the recognized potential of CFPD-based approaches as NAMs for simulating ACI, there is no first-principles-based CFPD model (i.e., a CFPD digital twin) that fully replicates the NGI system. To the best of our knowledge, the only CFD study of NGI primarily focused on stage-wise simulations or simplified geometries [13], thereby limiting its applicability to regulatory-relevant APSD prediction and comprehensive performance evaluation with physically realistic interactions between stages. Based on the literature review, there is a need to understand the spatiotemporal characteristics of NGI and to develop a CFPD-based *in silico model* of NGI to reduce the burden on laboratory experimental setups by shortening timelines and costs, thereby aiding early product development and approvals. Accordingly, this study developed and validated a CFPD-based virtual Next Generation Impactor (vNGI) as a fit-for-purpose in silico NAM for APSD prediction. The vNGI replicates the full NGI geometry (except for the MOC) and operating conditions (see **Fig. 1**), enabling detailed analysis of airflow structures and particle deposition across multiple stages, through systematic verification and validation against experimental data [9, 26, 27]. This work establishes the technical robustness, predictive capability, and regulatory relevance of the vNGI platform. Ultimately, this study advances NAMs in inhalation drug development by providing a scalable, mechanistically grounded tool to support bioequivalence assessment, device optimization, and early-stage product evaluation.

**Figure 1.**
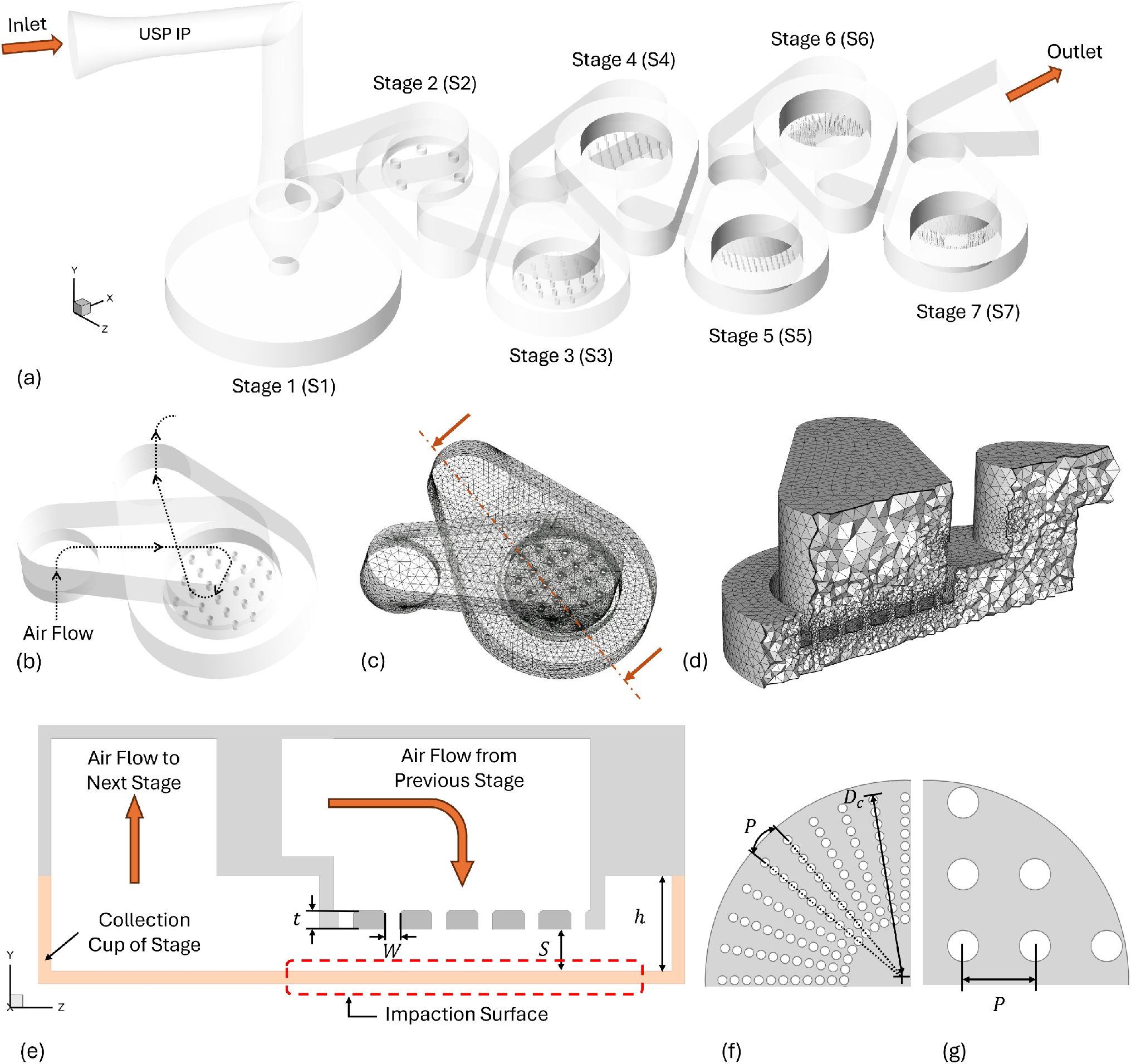
Geometry, mesh, and dimension details of the vNGI in this study: (a) Computer-aided design (CAD) geometry of the vNGI with the United States Pharmacopeia (USP) Induction Port (IP), (b) Filter nozzles of Stage 3 (S3) depicting air flow passage, (c) Surface mesh details of S3, (d) Cross-sectional view of tetrahedral-based volume mesh of S3, (e) 2D schematic diagram of nozzle and impaction plate, (f) Linear pitch dimensions used for Stage 3 (S3) to Stage 5 (S5), and (g) Angular pitch dimensions used for Stage 2 (S2), Stage 6 (S6) and Stage 7 (S7).

Also motivated by understudied topics raised by prior studies [28-33], the vNGI has been used in this study to investigate several key questions, i.e., the effect of air flow rate, mouth-throat geometry, and monodisperse vs. polydisperse conditions on particle transport and deposition in NGI, characterized by collection efficiency (*CE*) and deposition fractions (*DF*). Three airflow rates were considered, i.e., 15, 30, and 60 L/min. The simulated monodisperse particles covered a size range of 0.2 to 20.0 μm. For polydisperse particles, four log-normal distributions with different count mean diameters (CMDs) and count-based geometric standard deviations (GSDs) were used for injection [34]. In addition, three representative upper airway mouth-throat (MT) geometries were employed, including the United States Pharmacopeia (USP) Induction Port (IP), the Virginia Commonwealth University (VCU) Elliptic model (also called VE), and the VCU Realistic model (also called VR) (see **Fig. 2**).

**Figure 2.**
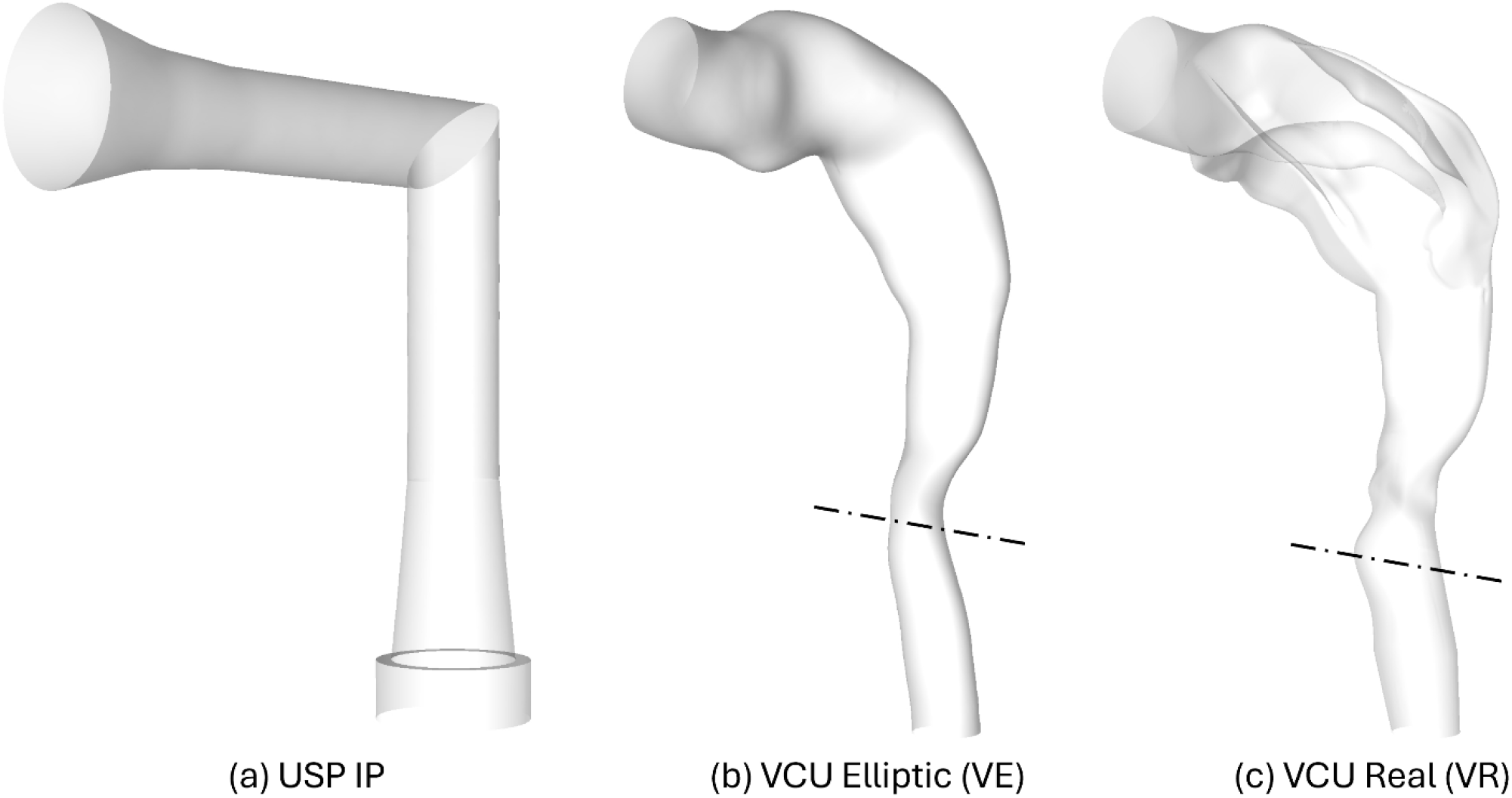
Computer-aided design (CAD) geometries of mouth-throat (MT) models employed in this study: (a) USP IP, (b) VCU Elliptic with sectional plane at glottis, and (c) VCU Real with sectional plane at glottis.

## 2. Methodology

### 2.1 Geometry and Mesh

#### 2.1.1 Geometry

The computer-aided design (CAD) geometry for the vNGI (see **Fig. 1**) was created in Ansys SpaceClaim 2024 R2 (Ansys Inc., Canonsburg, PA, USA). Specifically, the major geometric dimensions were derived from physical measurements of the cups, their spacings, their orientations relative to each other, the internal flow cavity, and the connecting flow channels (see **Table 1**). The critical dimensions, such as sieve sizes, the depth of the sieve inside the cups, and the pattern of sieves, were taken from the literature [9, 26], while the dimension of the induction port (IP) was taken from the U.S. Pharmacopeia [7]. Apart from the Reynolds number (*Re*) in the key design guidelines [26], the cross-flow parameter (*X*_*c*_) [35], and the nozzle-to-plate ratio (*S*/*W*) were kept well below the recommended 1.2 and 10 for a round nozzle while preparing the CAD geometry. Specifically, the cross-flow parameter *X*_*c*_ is defined as [35]

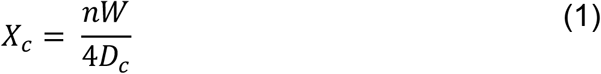

where *n, W* and *D*_*c*_ are the number of nozzles in a stage, nozzle diameter, and the diameter of the cluster of nozzles in a stage. The nozzle pitch (i.e., the angular or axial spacing between the nozzles) was appropriately adjusted to ensure the resultant *D*_*c*_ can lead to *X*_*c*_ values similar to those provided in [35], and the CAD dimensions are listed in **Table 1**. The other geometric dimensions listed in **Table 1** are throat length (*t*), the cup height (*h*), and volume (∀), the angular pitch (*P*) for S2, S6, and S7, as well as the axial pitch (*P*) for S3, S4, and S5. These dimensions are also marked in **Fig. 1**.

**Table 1.**
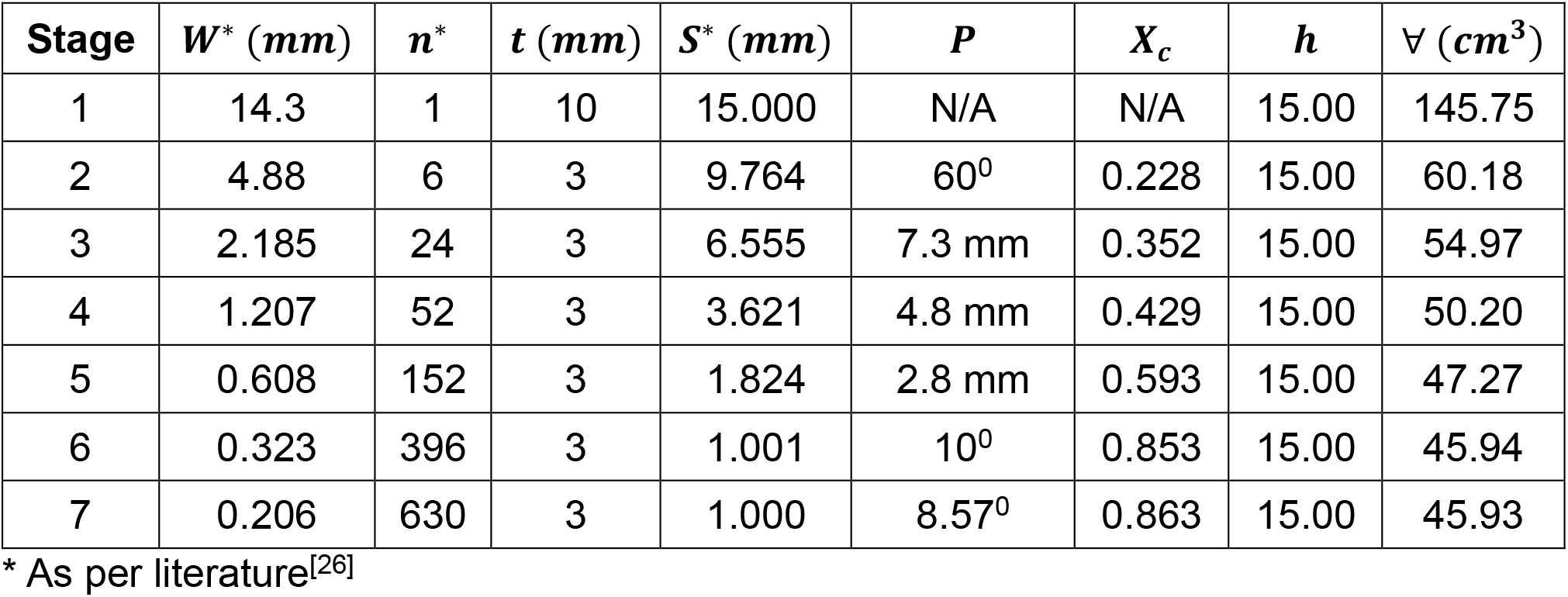
Geometric dimensions of the virtual Next Generation Impactor (vNGI)

| Stage | $W^*$ (mm) | $n^*$ | $t$ (mm) | $S^*$ (mm) | $P$ | $X_c$ | $h$ | $V$ (cm<sup>3</sup>) |
| --- | --- | --- | --- | --- | --- | --- | --- | --- |
| 1 | 14.3 | 1 | 10 | 15.000 | N/A | N/A | 15.00 | 145.75 |
| 2 | 4.88 | 6 | 3 | 9.764 | 60<sup>0</sup> | 0.228 | 15.00 | 60.18 |
| 3 | 2.185 | 24 | 3 | 6.555 | 7.3 mm | 0.352 | 15.00 | 54.97 |
| 4 | 1.207 | 52 | 3 | 3.621 | 4.8 mm | 0.429 | 15.00 | 50.20 |
| 5 | 0.608 | 152 | 3 | 1.824 | 2.8 mm | 0.593 | 15.00 | 47.27 |
| 6 | 0.323 | 396 | 3 | 1.001 | 10<sup>0</sup> | 0.853 | 15.00 | 45.94 |
| 7 | 0.206 | 630 | 3 | 1.000 | 8.57<sup>0</sup> | 0.863 | 15.00 | 45.93 |

\* As per literature<sup>[26]</sup>

To validate that the CAD geometry of the vNGI closely represents the physical NGI, the volume of the vNGI fluid domain was compared with the reported internal volume of the physical NGI [36]. The measured fluid-domain volume of the vNGI differs from the experimentally measured NGI volume by only 3.24%, indicating good geometric agreement between the vNGI CAD geometry and the physical NGI equipment.

It is worth noting that the micro-orifice collector (MOC) stage is not intended for measuring during the development of the physical NGI [26]. In a practical scenario to report formulation performance studies, the Mass Median Aerodynamic Diameter (*MMAD*) and Geometric Standard Deviation (*GSD*) is used, and the particle collection on the MOC stage is observed to be negligible in a broader range of OIDPs. [37-40]. From a computational standpoint, including the MOC stage would also substantially reduce simulation efficiency because it contains an extremely large number (i.e., 4032) of 70 µm nozzles, and the number of mesh elements required to represent the geometry in the simulation would increase exponentially, thereby increasing computational effort. Therefore, because of its limited practical significance and high computational cost, the MOC stage was omitted from the present study.

This study aims to extend the previous research [41], by incorporating vNGI with elliptic and realistic mouth-throat (MT) models created by researchers at Virginia Commonwealth University (VCU), and well studied [42-45] along with USP IP (see **Figs. 2 (a)-(c)**). Specifically, the diameters of the best circles fitting the mouth openings in the VCU Elliptic (VE) model (see **Fig. 2 (b))** and VCU Real (VR) model (see **Fig. 2 (c)**) are approximately 21.5 mm and 21.8 mm, respectively. They both yield an area of 366.43 mm^2^. Furthermore, the cross-sectional areas at the glottis for VCU Elliptic (VE) and VCU Real (VR) (see the dash-dot lines marked in **Figs. 2 (b) and (c)**) are 87.00 mm^2^ and 116.89 mm^2^, respectively. Additionally, the fluid-domain volumes of VCU Elliptic (VE) and VCU Real (VR) are 67.02 cm^3^ and 74.13 cm^3^.

#### 2.1.2 Mesh Independence Test

For the mesh independence test and to find the optimal mesh, three unstructured tetrahedral meshes with four near-wall prism layers (i.e., Coarse Mesh, Medium Mesh, and Fine Mesh) were generated for the vNGI (see **Figs. 1 (c) and (d)**) using Ansys Fluent Meshing 2024 R2 (Ansys Inc., Canonsburg, PA, USA). The growth ratio for prism layers and tetrahedral elements was kept at 1.12 and 1.18, respectively. The maximum skewness for the surface mesh was below 0.44. The orthogonal minimum quality was kept below 0.18. The average *y*^+^ was maintained across all wall surfaces, at approximately 1 or below, for all generated meshes.

The mesh independence test was conducted using CFD simulations with Coarse, Medium, and Fine Meshes, with cell counts of approximately 20.1 million, 24.5 million, and 29.5 million, respectively. To ensure that both mesh resolution and turbulence modeling were appropriately evaluated, the mesh independence study was conducted using three Reynolds-averaged Navier-Stokes (RANS) turbulence models at a steady-state volumetric flow rate of 30 L/min, i.e., k-ω Shear Stress Transport (SST) models, k-k_L_-ω, and the Generalized k-ω (GEKO).

These models were selected because they are appropriate for flows in inertial impactor systems and provide complementary capabilities in capturing near-wall and transitional flow behavior. Specifically, Previous studies on similar devices have used the standard k-ω model [21] and the k-k_L_-ω model [22]. In the present study, the k-ω SST model was selected as one of the candidates over the standard k-ω model because of its greater robustness and broader industrial acceptance, particularly for near-wall treatment [46, 47]. The k-k_L_-ω model was included because its additional transport equation for pre-transitional velocity fluctuations makes it well-suited for modeling laminar-to-turbulent transition [48, 49]. The GEKO model was also considered because it offers a good compromise between computational cost and the ability to capture local transition effects by tuning coefficients in the specific dissipation-rate equation [50, 51].

To evaluate mesh independence, the average velocity magnitude at 23,782 randomly distributed monitor points in the fluid domain was extracted and used to calculate the Average Percentage of Relative Difference (APRD) of velocity for each turbulence model across the tested meshes (see **Fig. 3**). The APRD was defined as

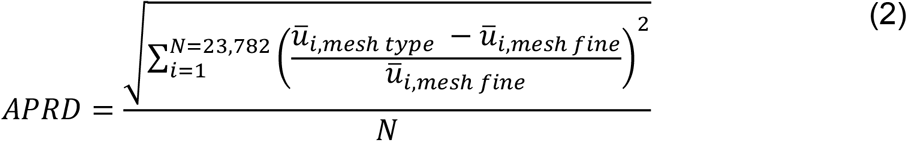

where *ū*_*i*_ is the time-averaged mean velocity magnitude at the monitor point *i*. The APRD of velocity decreased from the coarse mesh to the medium mesh, relative to the fine mesh, for all turbulence models. Similar trends were observed for the GEKO and k-k_L_-ω models, whereas the k-ω SST model exhibited a sharper reduction in error. Considering mesh sensitivity, turbulence-model performance, and mesh quality, the medium mesh with the GEKO model was selected for further study. This combination produced an ARPD of 0.65% compared with 0.62% for the coarse mesh, which is assumed acceptable given the balance between computational efficiency and accuracy. The meshing schemes, such as mesh element size and mesh element growth rate, were kept similar to those of USP IP for a bio-realistic MT model.

**Figure 3.**
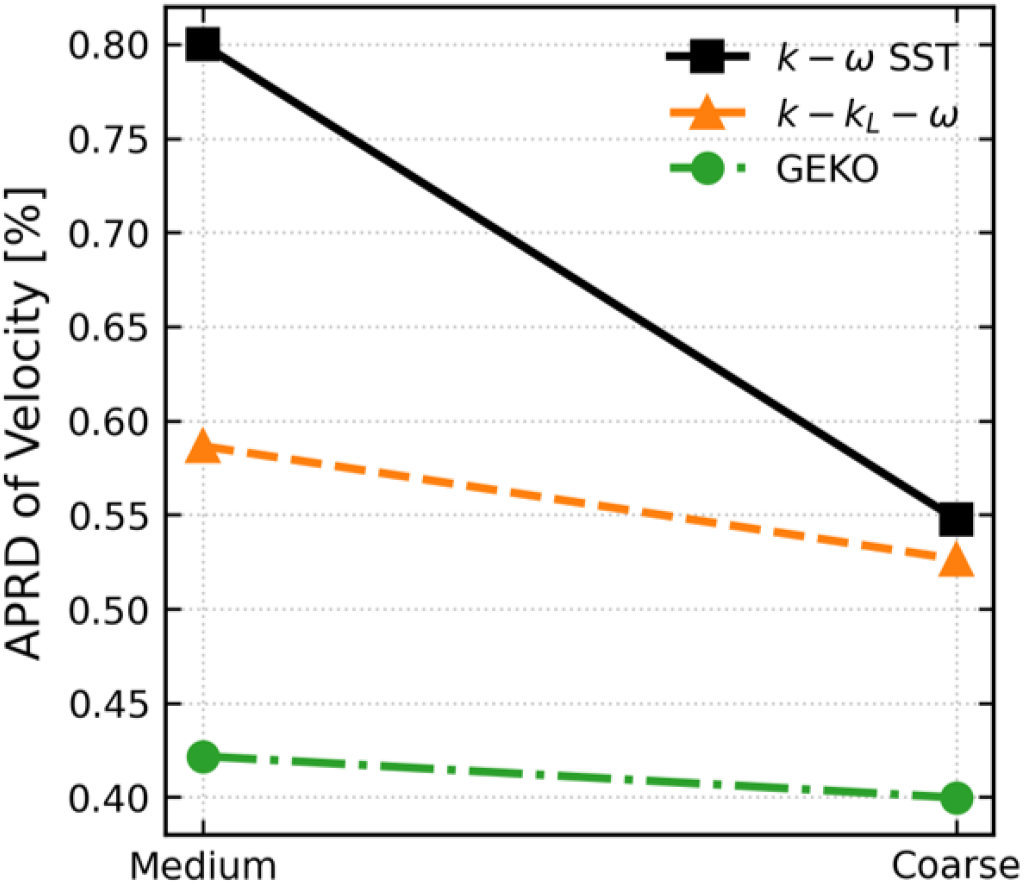
Comparison of airflow velocity error metrics at selected monitoring points in the vNGI CFD domain for different RANS turbulence models and mesh resolutions: Average percentage relative difference (APRD) of velocity magnitude as a function of mesh coarsening ratio relative to the fine mesh.

### 2.2 Governing Equations

#### 2.2.1 Continuous Phase (Airflow)

In practice, NGI is operated under steady-state flow conditions for all OIDPs testing at atmospheric pressure and room temperature, unless otherwise specified in the PSG, as with a spray metered dose inhaler [6]. Therefore, the airflow is assumed to be incompressible and isotropic with steady-state inlet flow rate conditions. The conservation laws of mass and momentum can be written in tensor form as

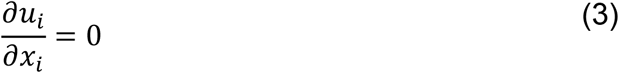

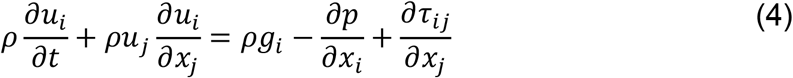

where vectors *u*_*i*_ and *x*_*i*_ represents fluid velocity and position, *S* is time, *g*_*i*_ is the gravitational acceleration, *ρ* is the air density, *p* is pressure, and *τ*_*ij*_ is the viscous stress tensor, which can be defined as [52]

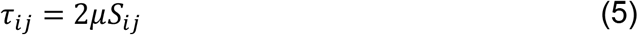

where *μ* is the air molecular viscosity and *S*_*ij*_ is the strain-rate tensor defined as

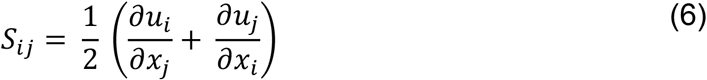

In the above equations, the flow velocity *u*_*i*_ and pressure *p* can be expressed as

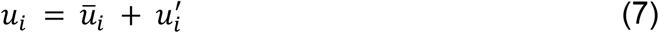

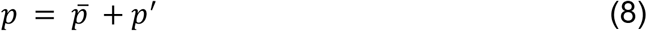

where *ū*_*i*_ and 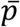 are the time-averaged mean velocity and pressure terms, while 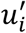 and *p*^′^ are the turbulence fluctuation terms. Equations (3) and (4) can be rewritten as

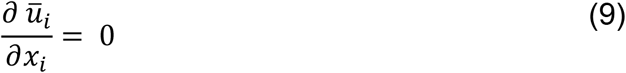

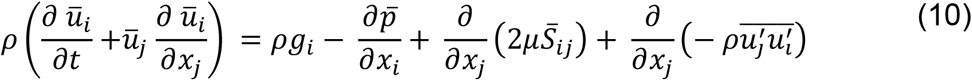

where

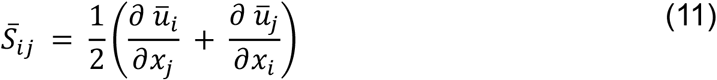

is the mean rate of strain tensor, and 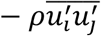 is the Reynolds stress. In the GEKO model for incompressible flow using the Boussinesq approximation [50, 51], 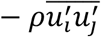 can be expressed as

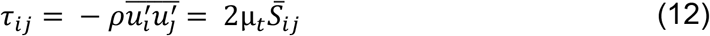

Formally, the stress tensor *τ*_*ij*_ contains an additional term 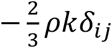. For Ansys Fluent Solver (Canonsburg, PA, USA), 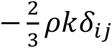 is presumed to be absorbed into the pressure gradient term [51]. *μ*_*t*_ is the eddy viscosity, which is expressed as

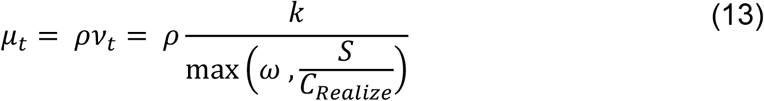

where *k* is the turbulent kinetic energy and *ω* is the specific dissipation rate. *ν*_*t*_ is the kinematic eddy viscosity and *S* can be defined as

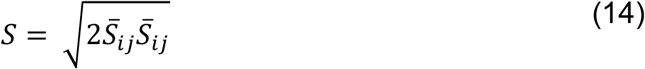

*C*_*Realize*_ is a GEKO main model coefficient with a value of 0.577. Accordingly, two transport equations for *k* and *ω* were added into the solving process, i.e.,

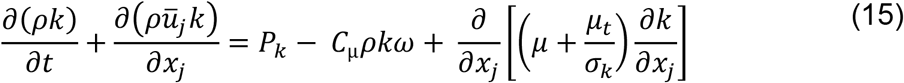

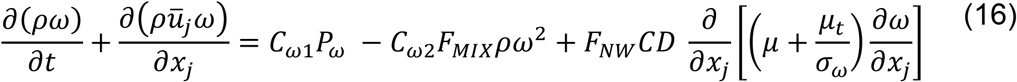

where *ū*_*j*_ is mean velocity tensor, *C*_*μ*_ and *C*_*ω*2_ are the GEKO main model coefficients with values 0.09 and 0.083, respectively. *σ*_*k*_ and *σ*_*ω*_ are turbulent Prandtl numbers for *κ* and *ω* respectively are defined as

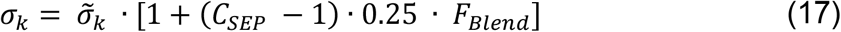

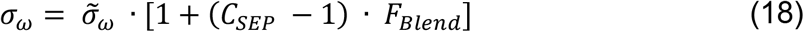

in Eqs. (17) and (18), 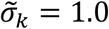 and 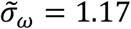 with the blending function *F*_*Blend*_ is defined as

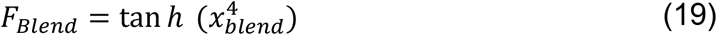

where *x*_*blend*_ is an intermediate nondimensional wall-proximity variable, which depends on an internal coefficient in GEKO, i.e., *CFb*_*Turb*_ = 2.0, *CFb*_*Lam*_ = 1.0, turbulence length scale *L*_*T*_, and wall distance *y*. Specifically, *x*_*blend*_ is defined as

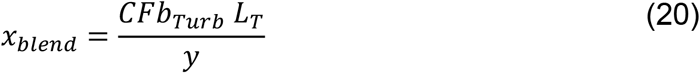

*L*_*T*_ can be given by

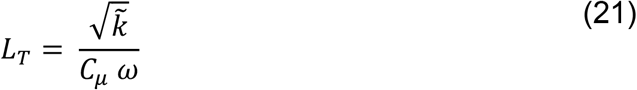

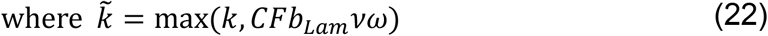

Furthermore, *P*_*k*_ and *P*_*ω*_ in Eqs. (15) and (16) are the net productions per unit dissipation of *k* and *ω*, respectively. *P*_*k*_ and *P*_*ω*_ can be defined as

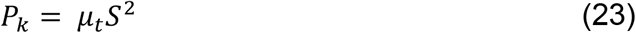

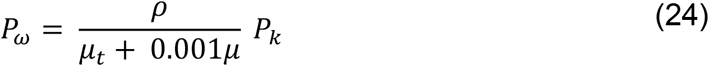

Additionally, *F*_*MIX*_ and *F*_*NW*_ in Eq. (16) are the independent formulations used to tune mixing-layer or boundary-layer flows and to preserve the law of the wall through near-wall damping [52], respectively. They are expressed as

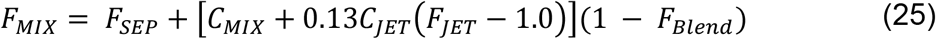

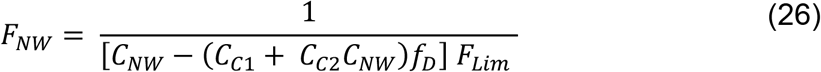

*CD* in Eq. (16) is a cross-diffusion term, which is defined as

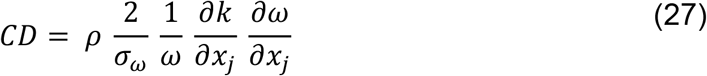

and *f*_*D*_ is a near-wall damping function in Eq. (26) defined as

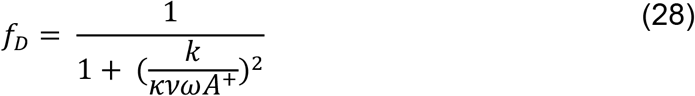

In Eq. (28), *κ* and *A*^+^are model constants associated with the von Kármán constant and Van Driest damping [53], respectively. Their default GEKO values are *κ* = 0.41 and *A*^+^ = 15.

In Eq. (26), *F*_*Lim*_ is a limiter function used for *CD*, which can be expressed as

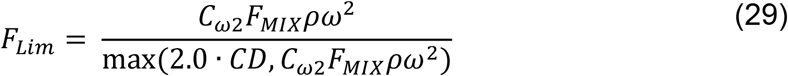

In Eq. (16), *C*_*ω*1_ is a variable used to keep the logarithmic near-wall behavior of the flow unaffected, and the formulation is expressed as

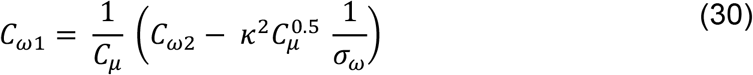

In Eqs. (16), (29) and (30), *C*_*ω*2_ = 0.083 is a model constant associated with the destruction term in the specific dissipation rate *ω* transport equation.

In Eq. (25), *F*_*JET*_ is the function used to tune jet flows is defined as

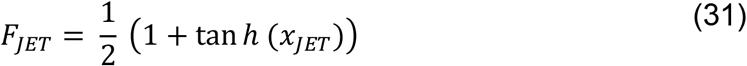

where *x*_*JET*_ is introduced as

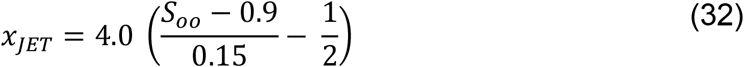

in which

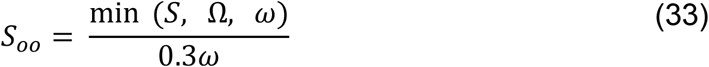

where Ω is the vorticity magnitude defined as

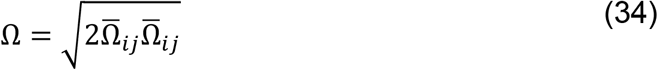

and 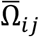 can be given as

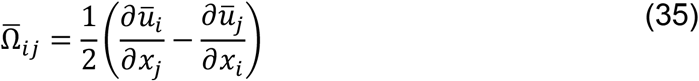

There are four free tunable coefficients, i.e., (1) C_SEP_ for flow optimization in separated flows, (2) C_NW_ for non-equilibrium near-wall regions, (3) C_MIX_ for mixing strength in free shear flows, and (4) C_JET_ for free shear-layer mixing. The default values used in the model are listed in **Table 2**. Based on these tunable coefficients, F_SEP_, F_NW_, F_MIX_, and F_JET_ are derived in the GEKO model.

**Table 2.** GEKO turbulence model parameters and their assigned values.

| **GEKO Parameter** | $C_{SEP}$ | $C_{NW}$ | $C_{MIX}$ | $C_{JET}$ |
| --- | --- | --- | --- | --- |
| **Assigned Value** | 1.75 | 0.5 | 0.0 | 1.0 |

#### 2.2.2 Discrete Phase (Particles)

The discrete phase, i.e., particles, was considered spherical, neglecting their rotational motion. It was assumed that the particle concentration was sufficiently dilute in the airflow field, making particle-particle interactions and the particle’s influence on the airflow field negligible. Therefore, the translational motion of the particle was solved by a one-way coupled Euler-Lagrange method [54]. Accordingly, the particle translational motion equation can be given as,

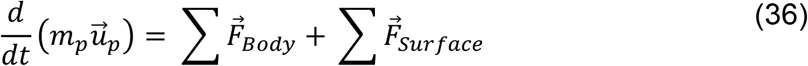

where 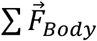 are body forces and 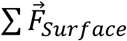 are surface forces acting on the particle. *m*_*p*_ and 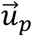 is the mass and velocity of the particle. Specifically, body forces and surface forces considered in this study are

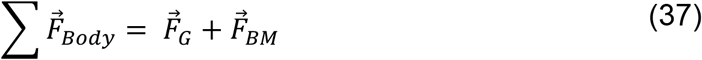

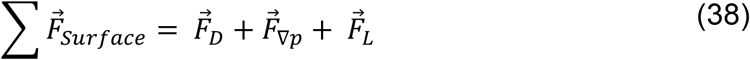

where 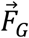 is the gravitational force, 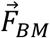 is the Brownian motion induced force [55, 56], 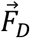 is the drag force, 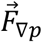 is the pressure gradient force, and 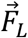 is the Saffman lift force [57].

The momentum exchange from the fluid phase to the aerodynamic particle diameter size above 1.0 *μm* was governed by the spherical drag law [58], whereas the aerodynamic particle diameters less than 1.0 *μm* were treated with Stokes-Cunningham law, in addition to Brownian motion [56]. The stochastic tracking was enabled with a random eddy lifetime. The higher-order tracking scheme employed was the Runge-Kutta method. The particle density was assigned as the standard aerodynamic density *ρ*_*ae*_ = 1000 *kg*/*m*^3^. The density ratio of fluid to particle density was considered negligible; the other forces exerted on the particles were not considered.

The Cunningham slip correction factor (*C*_*c*_) [59] was used for calculating 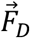 for all the particles of diameter size below 1.0 *μm*, expressed as [26]

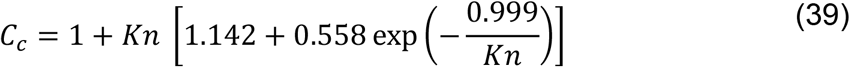

where *Kn* is the Knudsen number, which can be defined as [26]

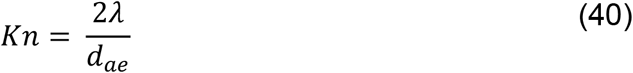

in which *λ* is the air mean free path, and *d*_*ae*_ is the aerodynamic diameter of the particle.

#### 2.2.3 Particle Collection Efficiency

For monodispersed particles, the particle collection efficiency (*CE*) of a specific zone (e.g., a stage) in this study can be defined as [9]

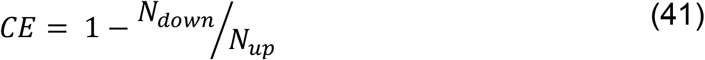

where *N*_*down*_ is the number of particles deposited downstream of the zone, and *N*_*up*_ is the number of particles entering the zone from upstream.

#### 2.2.4 Particle Deposition Fraction

The particle deposition fraction (*DF*) is adopted from the literature [54] in percentage, as the total mass of particles deposited in a specific region divided by the total mass of particles injected at the mouth-throat inlet can be defined as,

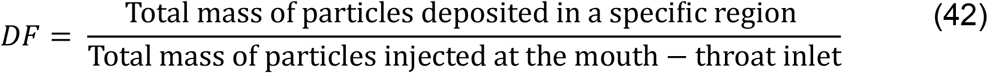

### 2.3 Material Properties

#### 2.3.1 Air

The air flow field was considered as an incompressible fluid at an ambient temperature of 300*K* and a pressure of 1 *atm*, having a density *ρ* as 1.225 *kg*/*m*^3^ with viscosity *μ* as 1.7894 × 10^−5^ *kg*/*m* · *s*.

#### 2.3.2 Particles

The intent of the CFPD simulations using vNGI in this study was to visualize and analyze the spatiotemporal distributions of air-particle multiphase flow. Therefore, the particles were considered solid, spherical, and chemically nonreactive with the airflow and the vNGI walls. The particle density *ρ*_*p*_ was equal to the standard aerodynamic density *ρ*_*ae*_ = 1000 *kg*/*m*^3^.

The monodisperse particles simulated were in a particle diameter range of 0.20 *μm* ≤ *d*_*p*_ = *d*_*ae*_ ≤ 20.0 *μm*. For polydisperse particles, four log-normal distributions were considered for injection [34]. The particle number distribution 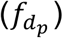 (also called particle number frequency) as a function of particle diameter was calculated by

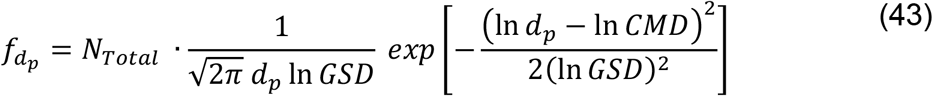

where *CMD* is the count median diameter, *GSD* is the associated geometric standard deviation, and *N*_*Total*_ is the total number of injected particles. Details of the four polydisperse particle size distributions injected (i.e., Cases 1-4) can be found in **Fig. 4** and **Table 3**.

**Table 3.** Prescribed vs. achieved lognormal particle size distribution parameters for polydisperse aerosol injection in vNGI simulations: input *CMD*/*GSD* and actually injected count-based *CMD*/*GSD*, with corresponding mass-based *MMD* and *GSD*.

| **Case** | **Input**<br>***CMD***<br>**($\mu m$)** | **Input**<br>**Count**<br>***GSD*** | **Actual**<br>***CMD* ($\mu m$)** | **Actual**<br>**Count *GSD*** | **Actual**<br>***MMD* ($\mu m$)** | **Actually**<br>**Mass *GSD*** |
| --- | --- | --- | --- | --- | --- | --- |
| **Case 1** | 1.0 | 1.5 | 0.9519 | 1.5436 | 1.5808 | 1.5142 |
| **Case 2** | 1.0 | 1.8 | 0.9516 | 1.8680 | 2.6482 | 1.7459 |
| **Case 3** | 2.5 | 1.5 | 2.4496 | 1.5145 | 4.0112 | 1.4947 |
| **Case 4** | 2.5 | 1.8 | 2.4348 | 1.8102 | 5.6905 | 1.5518 |

**Figure 4.**
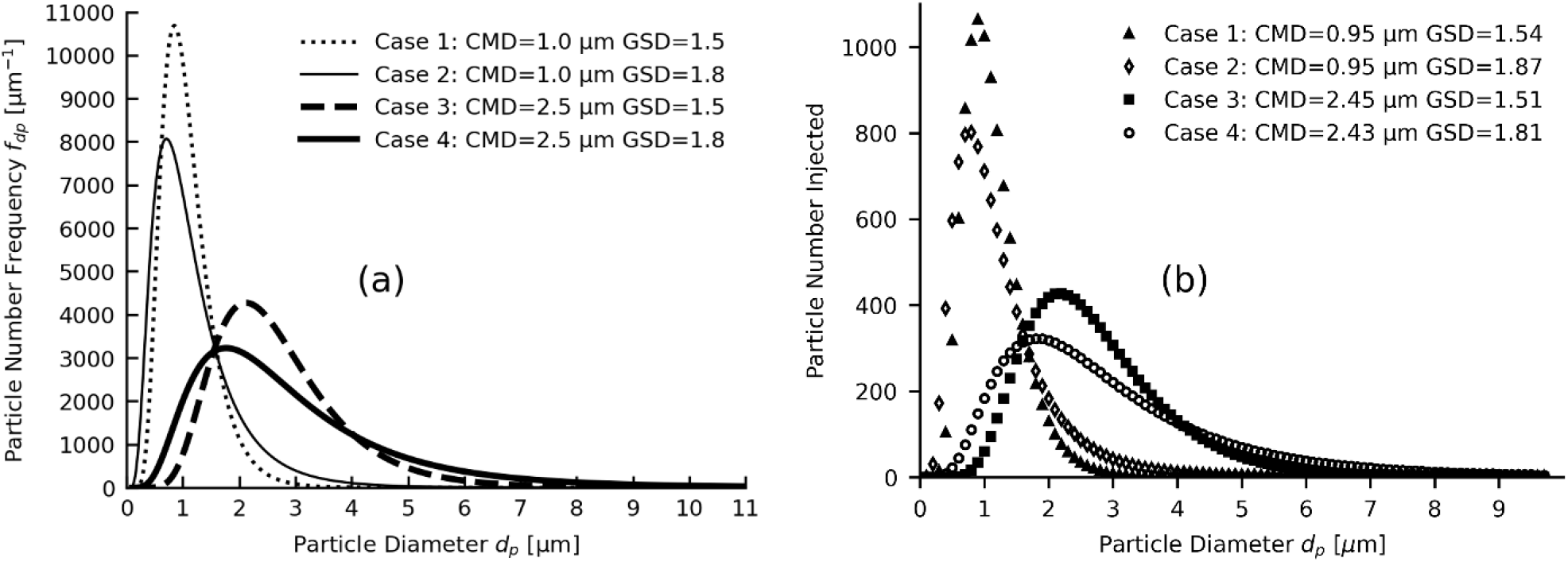
Log-normal particle size distribution for polydisperse particle injection with varying count median diameters (*CMDs*) and associated geometric standard deviations (*GSDs*) in this study (see Table 3) : (a) Continuous particle frequency distributions (see Eq.(43)), and (b) Discrete particle number injections implemented CFPD simulations corresponding to the specified *CMD* and *GSD* combinations.

### 2.4 Initial and Boundary Conditions

For airflow, constant velocity inlet conditions corresponding to three flow rates, i.e., 15 L/min, 30 L/min, and 60 L/min at the mouth inlets of all three mouth-throat geometries (see **Fig. 2**) were applied. Zero-gauge pressure outlets were implemented at the exit of vNGI. All the walls of vNGI were considered no-slip for velocity. The walls of the domain were considered to have a smooth surface with zero surface roughness.

For particles, monodisperse and polydisperse particles were individually injected at the mouth-throat model inlets for the calculation of individual *CE* for monodispersed and *DF* for polydisperse particle-laden flow. All particles were injected at random locations within the three M-T model inlets via file injection. For every monodispersed particle diameter size, 10,000 particles were finalized based on a particle count independence test conducted on a discrete particle size range over the particle number counts of 10,000, 50,000, and 100,000 particles. No significant change in *CE* was observed after varying the particle count. The particle time step independence study was also conducted, and the particle time step was finalized at 1.0e-6 s, with the fluid flow time step at 0.01 s. Based on the outcomes from monodisperse particles, approximately 10,000 polydisperse particles for Cases 1-4 were injected. The fate of particles hitting the wall surface was considered, as particles were assumed to deposit except at the filter walls, where they may re-entrain due to very high shear flow. Therefore, a reflective boundary condition was implemented for particles hitting the filter walls, since deposited particles tend to detach from them [21]. The fate of particles reaching the vNGI outlet was considered as deposited. The particles deposited at the outlet of the vNGI were used as a pseudo-analysis of particle deposition at MOC, since the remaining particles from S7 are most likely to deposit at MOC per NGI design considerations [9, 26].

### 2.5 Numerical Setup

Initially, the steady-state airflow simulations were conducted using Ansys Fluent 2024 R2 (Ansys Inc., Canonsburg, PA) on a local Dell Precision T7960 workstation (Intel® Xeon® Processor w9-3575X (2.21 GHz), 8 cores, and 256 GB RAM). A semi-implicit pressure-linked equation (SIMPLE) algorithm was used for pressure-velocity coupling with the Green-Gauss node-based scheme to calculate the node-based gradients. The second-order scheme was employed for pressure discretization. The second-order upwind scheme was used to discretize the momentum and turbulent kinetic energy equations. The residual convergence criteria were kept below 1.0e-5. The secondary convergence criteria were based on measurements of Reynolds number (*Re*) at locations near each filter exit (see **Fig. 5(a)**). The solution is considered converged when the reduction rate of the *Re*’*s* residual is sufficiently low. The steady-state airflow fields were loaded in the one-way coupled Euler-Lagrange method (i.e., DPM) to predict transient particle transport and deposition in the vNGI. The DPM simulations were advanced until fewer than 1.0% of the injected particles remained suspended within the computational domain, or until a physical time of 13 s was reached. The maximum simulation duration was determined based on computational efficiency and practical considerations [37]. All discrete phase simulations were performed on the Pete high-performance computing (HPC) cluster at Oklahoma State University (OSU).

**Figure 5.**
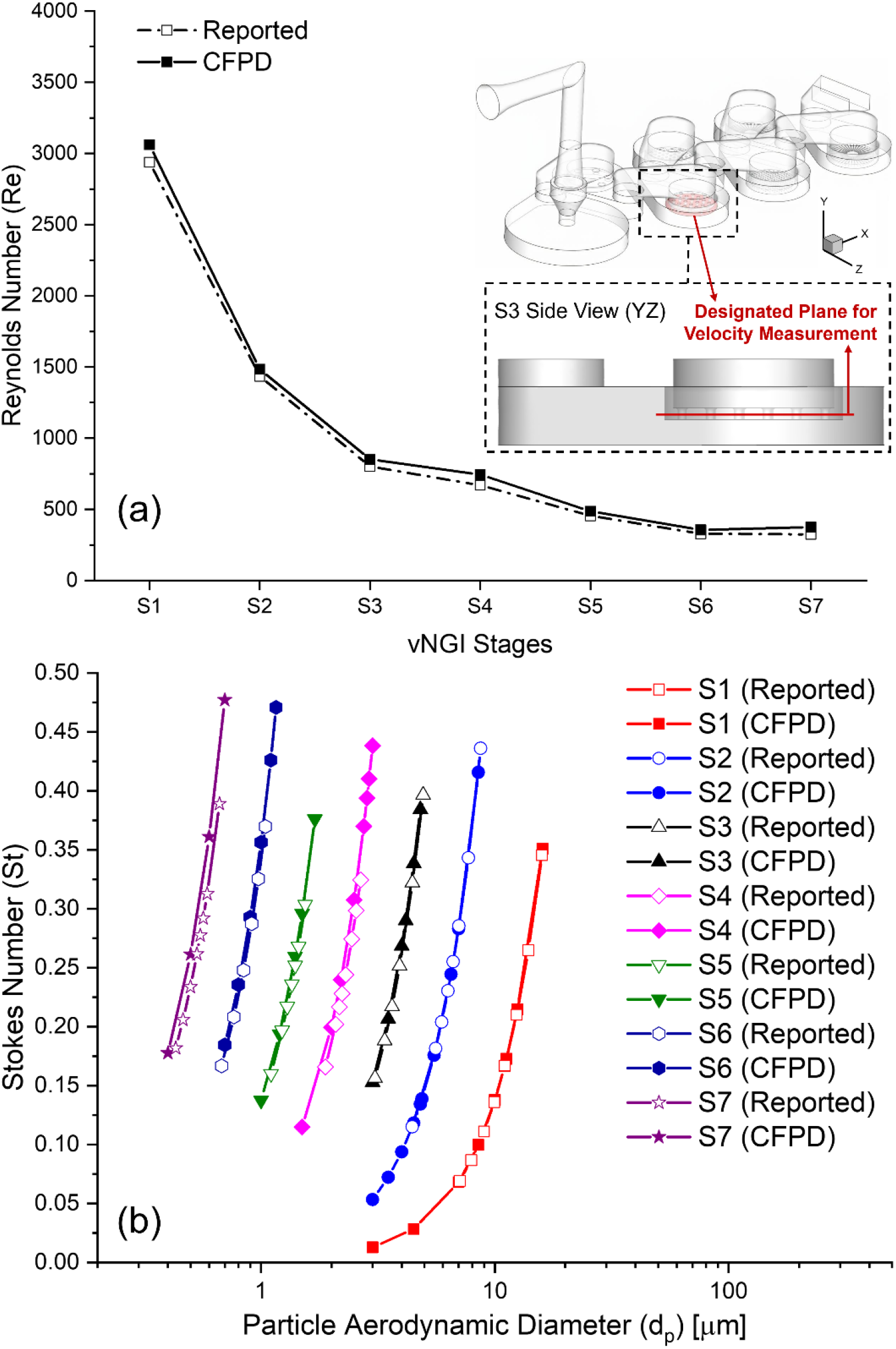
Model validation of CFPD predictions against reported data ^[26]^ at a steady inhalation flow rate (30 L/min): (a) Reynolds numbers (*Re*) across vNGI stages with indicated velocity sampling locations and (b) Stokes number (*St*) at different particle diameters

### 2.6 Model Validations

#### 2.6.1 Airflow Field

The airflow field is one of the important aspects [60, 61] to define a sharp cutoff of the stage, which could be validated by comparing predicted *Re* and *St* with the presented work in the literature [26, 27]. The expression used in literature for analytically calculating the *Re*_*Reported*_ can be given as,

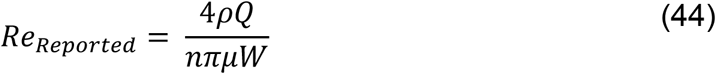

where *Q* is the volumetric flow rate at the inlet.

In this study, the predefined air properties, such as density (1.225 *kg*/*m*^3^) and viscosity (1.7894 × 10^−5^ *kg*/*m* · *s*), were used. Using these properties, analytically calculated *Re*_*Reported*_ using Eq. (44) was found about 3.7 % higher than reported in the literature [26] for all flow rates. The difference could be due to variations in the air properties considered. Therefore, the calculation difference was supposed to be carried forward in *St* and other allied computations. In an existing study [9, 26], the *St* for impactors was calculated based on the global variable as the volumetric flow rate at the inlet using a modified *St* for round jets [26, 62, 63], which is defined as

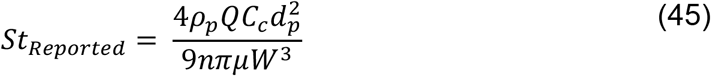

This study assumed that the local velocity at the exit of the nozzle plays a significant role in determining the *St* and could provide a better understanding of particle deposition. Therefore, the expressions in this study to calculate CFPD predicted *St* and *Re* using velocity (*V*_0_) at the exit of the nozzle are defined as

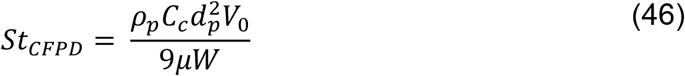

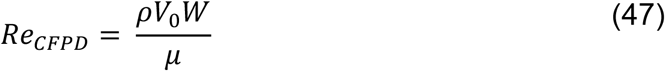

As shown in **Fig. 5(a)**, *Re*_*CFPD*_ values predicted using CFPD-based vNGI at 30 L/min were found in close agreement with the literature data [26] which is calculated analytically using Eq. (44) at all stages. The difference between the CFPD predicted *Re* and reported values (see **Table 4**) were quantified using a relative difference percentage of *Re* (*δRe*_*rel*_), which is defined as

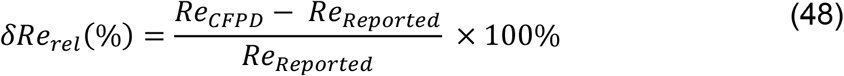

where *Re*_*CFPD*_ is the CFPD-predicted *Re*, and *Re*_*Reported*_ is the reported value [26, 27]. Across all stages, *Re*_*CFPD*_ was found to be slightly higher than *Re*_*Reported*_. In case of 30 L/min flow rate, *δRe*_*rel*_ for S1 and S2 is 4.2% and 3.4%, whereas for S3, S5, and S6 *δRe*_*rel*_ are 6.2%, 7.0%, and 8.5% higher, respectively. The S7 showed the highest relative difference of 15.5% among all.

**Table 4.** Comparison of the CFPD predicted *Re* with data in open literature [26, 27].

| Flow Rate [L/min] | Stage No. | $Re_{Reported}$ | $Re_{CFPD}$ | $\delta Re_{rel}$ (%) |
| --- | --- | --- | --- | --- |
| 15 | S1 | 1482 | 1534.21 | 3.52 |
|  | S2 | 724 | 742.38 | 2.54 |
|  | S3 | 404 | 427.32 | 5.77 |
|  | S4 | 338 | 375.40 | 11.06 |
|  | S5 | 230 | 243.20 | 5.74 |
|  | S6 | 167 | 178.46 | 6.86 |
|  | S7 | 166 | 188.12 | 13.33 |
| 30 | S1 | 2938 | 3061.45 | 4.20 |
|  | S2 | 1435 | 1484.35 | 3.44 |
|  | S3 | 801 | 851.05 | 6.25 |
|  | S4 | 669 | 744.65 | 11.31 |
|  | S5 | 455 | 486.93 | 7.02 |
|  | S6 | 328 | 355.93 | 8.52 |
|  | S7 | 324 | 374.23 | 15.50 |
| 60 | S1 | 5876 | 6142.87 | 4.54 |
|  | S2 | 2870 | 2981.92 | 3.90 |
|  | S3 | 1602 | 1700.13 | 6.13 |
|  | S4 | 1339 | 1481.05 | 10.61 |
|  | S5 | 909 | 980.40 | 7.86 |
|  | S6 | 657 | 712.39 | 8.43 |
|  | S7 | 647 | 747.03 | 15.46 |

In literature, analytically or experimentally determined *St* are not presented [9, 26, 27]. Therefore, known air properties which are used for prediction of *Re* were used as mentioned earlier to calculate *St* at each stage using Eq. (45) and henceforth these values should be considered as reported values in literature [9, 26, 27]. The comparison of predicted *St* with the reported values was on the high side, but in excellent agreement for all stages (see **Fig. 5(b)**) at a 30 L/min. Hence, based on the comparison of *Re* and *St*, the capability of the CFPD model to predict airflow field in NGI is considered validated.

Similarly, when the CFPD predicted *Re* and *St* were compared with the reported data at 15 L/min [27] and 60 L/min [26], the comparability for *Re* and *St* was found to be indifferent to the trend of 30 L/min (see **Table 4**). In the case of 15 L/min, CFPD predicted *Re* for S1-S3 was found between 2.5% to 5.7% higher, whereas for S5 it was 5.7% and for S6 it was 6.8% higher. The relative error for S4 and S7 was found to be 11.0% and 13.3% higher, respectively, with the reported data [27] (see **Fig. 6(a)**). In case of 60 L/min, the CFPD predicted *Re* for S1 and S2 was 4.5% and 3.9%, were relatively higher than the reported data [26]. The other stages also showed relatively higher CFPD predicted *Re*, such as for S3-S7, it was found 6.1%, 10.6%, 7.8%, 8.4%, and 15.4% higher, respectively (see **Fig. 6(c)**). In comparison, the trend in CFPD predicted *St* and reported *St* in the literature can be referred from **Fig. 6 (b) and (d)**, which was found in excellent agreement for both the 15 L/min and 60 L/min steady inhalation flow rates. The predicted *St* for S7 was found to show the highest 6% deviation, whereas for all the other stages it was within 1%-5% for both flow rates. This comparative study of different flow rates further strengthened the confidence in the CFPD-based vNGI model.

**Figure 6.**
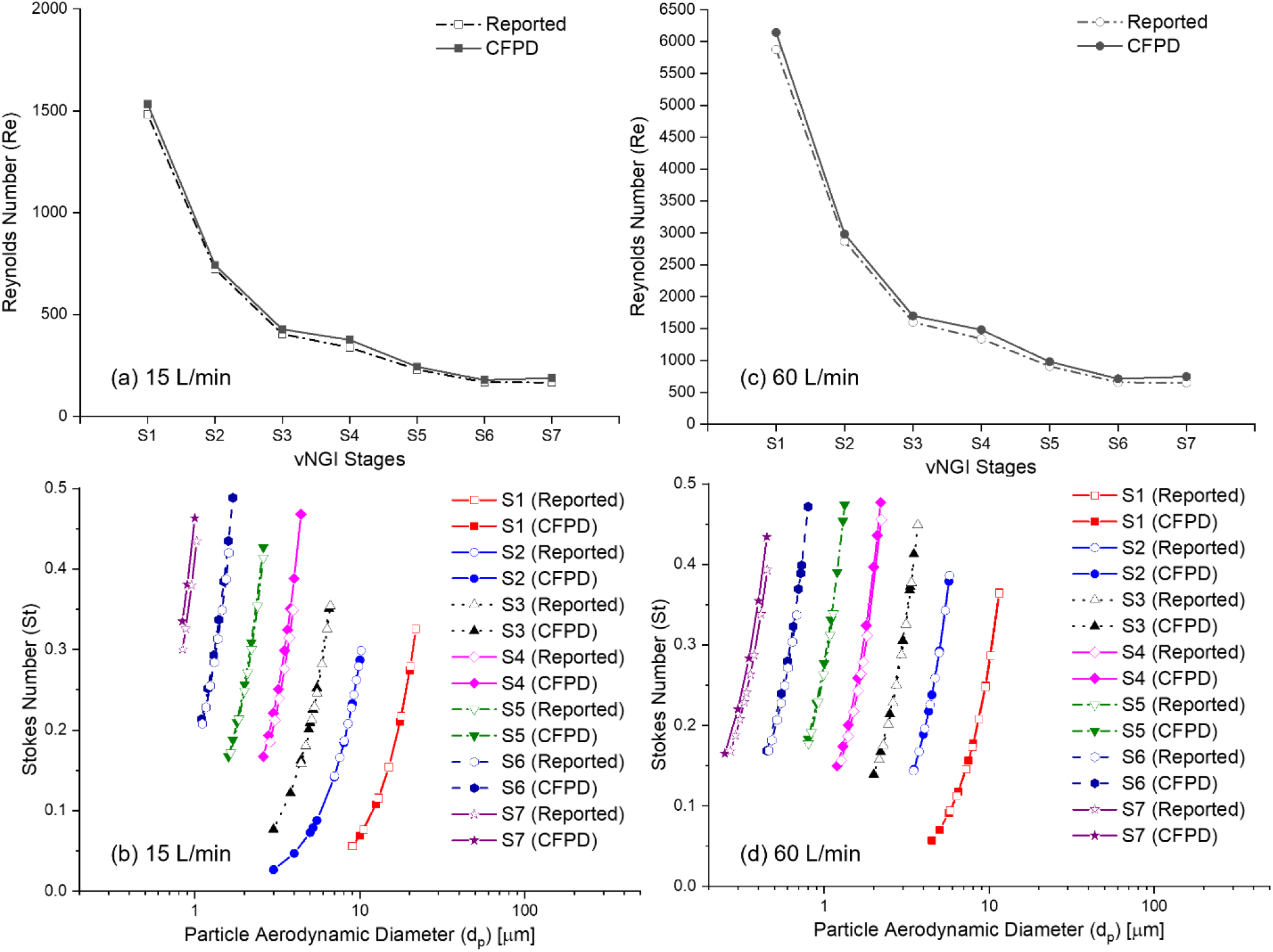
Comparison of CFPD predicted data with the reported values ^[26, 27]^ at each stage at different steady flow rates: (a) Reynolds numbers (*Re*) across vNGI stages at a steady inhalation flow rate (15 L/min), (b) Stokes number (*St*) with different particle diameters at a steady inhalation flow rate (15 L/min), (c) Reynolds numbers (*Re*) across vNGI stages at a steady inhalation flow rate (60 L/min), and (d) Stokes number (*St*) with different particle diameters at a steady inhalation flow rate (60 L/min)

#### 2.6.2 Particle Deposition

The validation of the discrete phase model was conducted for 30 L/min only by comparing the CFPD predicted cut-off diameter (*d*_50_) (see **Fig. 7(a)**) and *CE* (see **Fig. 7(b)**) in each stage with the experimental data [9]. *d*_50_ was calculated by implementing the weighted linear regression method [64], similar to what has been employed in the experimental investigation [9]. The data points used for the calculation of *d*_50_ were considered between *d*_16_ and *d*_84_, closer to *d*_50_ for calculation by interpolation to preserve the maximum linearity in the sharper regions of the *CE*.

**Figure 7.**
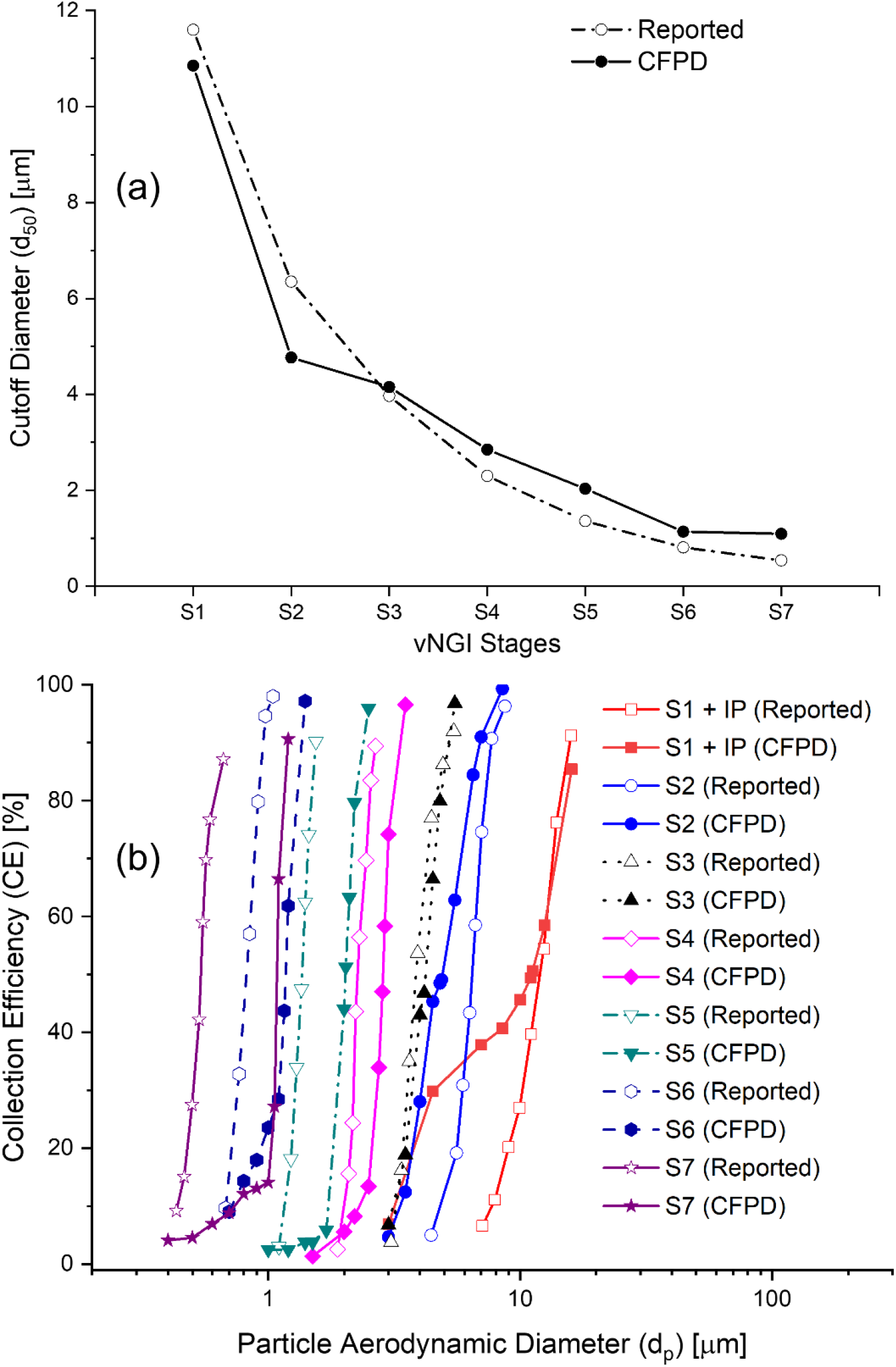
Comparison of CFPD predicted data with the reported values ^[9]^ at each stage for 30 L/min steady flow rate (a) Particle cut-off diameter (d_50) prediction and (b) Particle collection efficiency (CE) prediction

For the *CE* comparisons shown in **Fig. 7(b)**, monodisperse particles of different diameters were injected into the resolved airflow field at 30 L/min, and *CE* was calculated for each vNGI stage. As shown in **Fig. 7(b)**, the predicted *CE* curves exhibited the expected sigmoidal trend (also mentioned as S shape). They were generally found to be consistent with the experimental measurements [9]. The lower the inclination at the vertical part near the 50% of a *CE* curves, the higher chances of achieving the sharp diameter cutoff (*d*_50_) at that stage. Agreement was strongest for S3, while S1, S2, S4, and S5 showed reasonable agreement. In contrast, the predicted *CE* curves for S6 and S7 nearly overlapped, resulting in very similar predicted cut-off diameters (*d*_50_) (see **Fig. 7(a)**). The relative difference between the predicted cut-off diameters of S6 and S7 was less than 4%, whereas the experiment [9] reports a difference of about 33% between the stages. The overlap of the CFPD predicted *CE* curves for S6 and S7 is likely attributable to a limitation of the DPM approach. Because DPM treats particles as point masses, the finite particle size is not fully represented during transport through the stages. Consequently, particles that are large enough to be restricted by the small nozzle geometry in the real NGI may still penetrate stage 7 in the simulation. This can lead to overprediction of particle penetration into the later stages and, in turn, unrealistically similar deposition efficiencies and cut-off behavior for stages 6 and 7. Thus, the overlap between these two stages is more likely a modeling artifact than a physically realistic deposition pattern. Furthermore, particle-particle interactions were also neglected in this study. Including such interactions could increase particle deposition in upstream NGI stages due to particle-particle agglomeration, this would shift the *CE* to the right. Additionally, all particles impacting non-collection walls were assumed to deposit completely. This treatment assumes fully adherent wall surfaces and may increase predicted interstage wall losses. A large fraction of the deposited particles was found upstream of the filters. Since no detailed NGI studies on interstage wall losses were identified, complete deposition was assumed instead of introducing uncertain partial restitution or wall rebound behavior. Experimental uncertainty may also contribute to the discrepancies. Possible sources include failure to maintain the exact pressure drop across the filter nozzle [65], neglect of particle cloud settling effects in the experimental setup [62], and other setup modifications required to accommodate calibration instruments [9].

For the comparisons of cutoff diameters (*d*_50_) shown in **Fig. 7(a)** with 30 L/min, the overall trend in the predicted cutoff diameters was consistent with the experimental data [9]. The first two stages (i.e., S1 and S2) underpredicted *d*_50_, whereas the downstream stages (i.e., S3-S7) overpredicted it. S3 served as the transition point between underprediction and overprediction and showed the smallest relative error (i.e., 4.64%). From S4 to S7, the relative errors increased progressively to 24%, 50%, 41%, and 103%, respectively. For stages 1 and 2, the relative errors in cut-off diameter were 6.5% and 25%, respectively.

Additionally, studies have reported that for a sharp cutoff, *Re* should be in the range of 500 to 3000 [34, 61, 66, 67]. However, the last two stages (i.e., S6 and S7) are designed to achieve *Re* approximately equal to 320, which is lower than the recommended, and both have almost similar *Re* (328 and 324 for S6 and S7, respectively at 30 L/min). In the impactor studies, it has been reported that for *Re* < 500, Poor cutoffs were observed. It could be due to the thick viscous boundary layer in the jet of the impactor nozzle [62, 63]. This could be because particles with low velocity gain less momentum, leading to lower deposition, and because particles traveling in the boundary layer have lower velocity than those in the central part of the jet. Similarly, for higher *S*/*W*, the intensity of the sharp cut-off reduces on the *CE* curve with respect to the 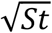 [63, 66], where *S*/*W* is 3.1 and 4.9 for S6 and S7, respectively.

Hence, the *CE* prediction using the vNGI in S6 and S7 is considered non-ideal. Despite such limitations, the overall validation results indicate that the CFPD-based NGI still captures the stage-wise deposition characteristics very well in S1 to S5. In inhalation product development, deposition across multiple stages is often grouped to evaluate APSD using NGI [68], which reduces the influence of stage-specific discrepancies in *CE* and *d*_50_ between the vNGI predictions and the experimental data [9]. Therefore, although the current vNGI model has limitations in predicting depositions in S6 and S7, the validations shown in **Fig. 7** support its potential for further research. In particular, the CFPD-based vNGI remains valuable for investigating flow and particle transport phenomena that are difficult or impossible to study experimentally. Considering the inherent limitations of computational modeling and the lack of sufficiently detailed experimental data for complete cross-validation, the vNGI model provides a useful and worthwhile tool for further analysis.

## 3. Results and Discussion

### 3.1 Airflow Field

Beyond 30 L/min simulations for validation (see **Section 2.6.1**), the vNGI model was further tested for its predictive capabilities at steady-state ventilation flow rates of 15 L/min and 60 L/min. Similar to 30 L/min, the steady state airflow field simulations was conducted for other flow rates, and the predicted *Re* and *St* at all flow rates were compared with the reported data [9, 27] (see **Figs. 5 and 6**).

Although the NGI is designed for keeping stage-wise *Re* between 500 and 3000 with *St*_50_ about 0.245 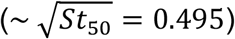 and limiting *S*/*W* within 1-10 [9, 26], limiting the discussion with non-dimensional important parameters such as *Re* and *St* may not be sufficient to understand the entire flow dynamics of the intricate NGI. The major structure of the design consists of lateral converging and diverging cross-sectional areas (see **Fig. 1**), causing the flow to experience frequent favorable and adverse pressure drops, respectively, as it travels from one stage to another. The purpose of this could be to decelerate the airflow before reaching the nozzle filters by divergent cross section, so that the particles get sufficient time to accelerate again at the filter nozzle. Then, after the particle impaction at the stage, the flow field is accelerated by a converging cross-sectional area. In this way, the particles are neither decelerated enough to settle nor accelerated enough to avoid impacting these interstage areas. The first stage can be considered an ideal single-round jet inertial impactor, which is consistent with theoretical understanding. Still, other stages are a type of multi-nozzle round jet inertial impactor and have relatively uniform flow distribution among the filter nozzles within the stage caused by upstream flow conditions (see **Figs. 8 and 9**) at all three flow rates. The pressure drops through the filters increase from S1 to S7 (see **Fig. 10** for a comparison between S3 and S6 as an example). Specifically, at S3, the pressure drops are approximately 10 Pa, 40 Pa, and 159 Pa at 15, 30, and 60 L/min, respectively. In contrast, the pressure drop increases by 228 Pa, 618 Pa, and 1870 Pa as the flow rate increases.

**Figure 8.**
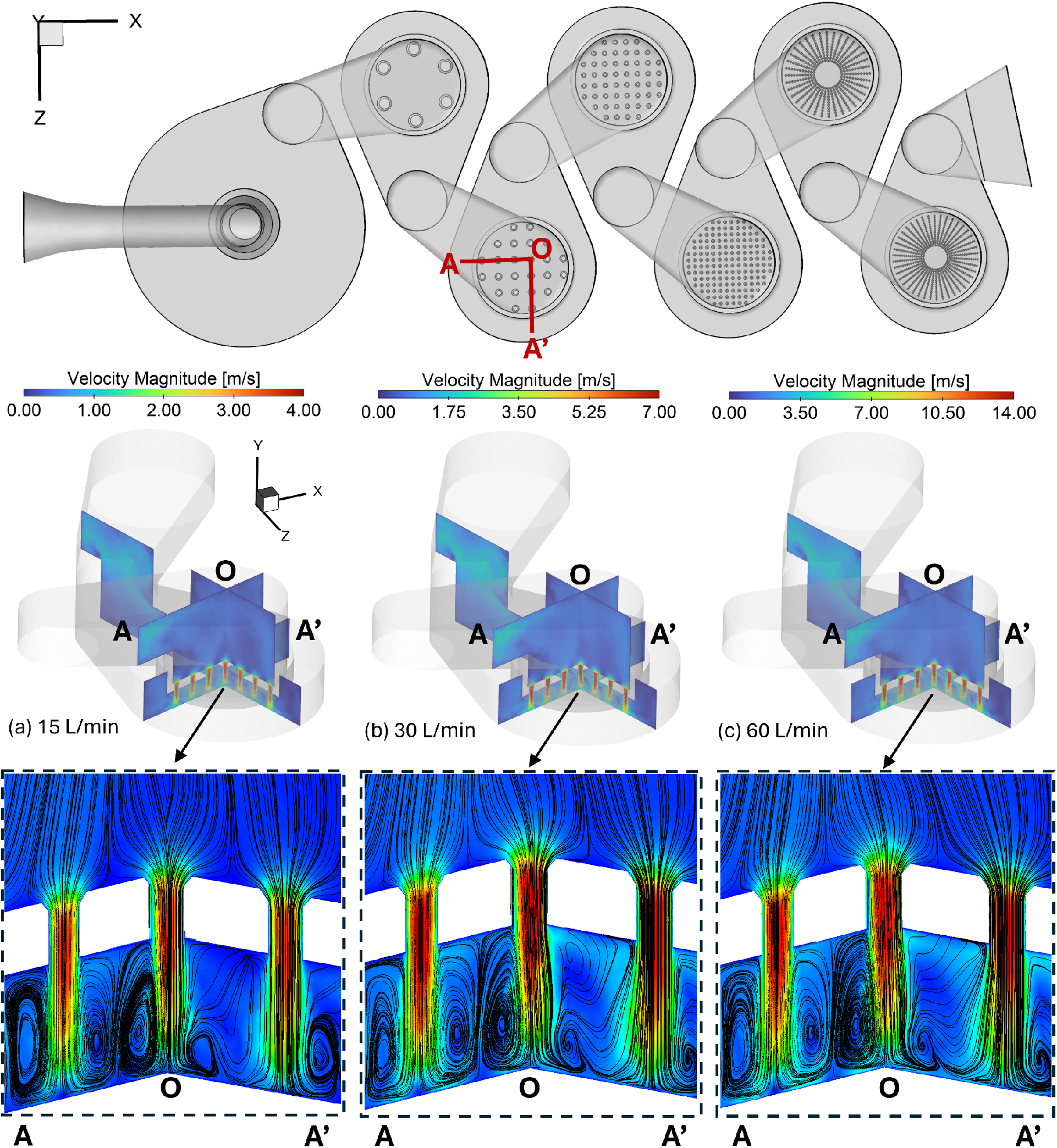
Velocity magnitude contours and streamlines at the filter of Stage 3 (S3) at three different flow rates: (a) 15 L/min, (b) 30 L/min, and (c) 60 L/min

**Figure 9.**
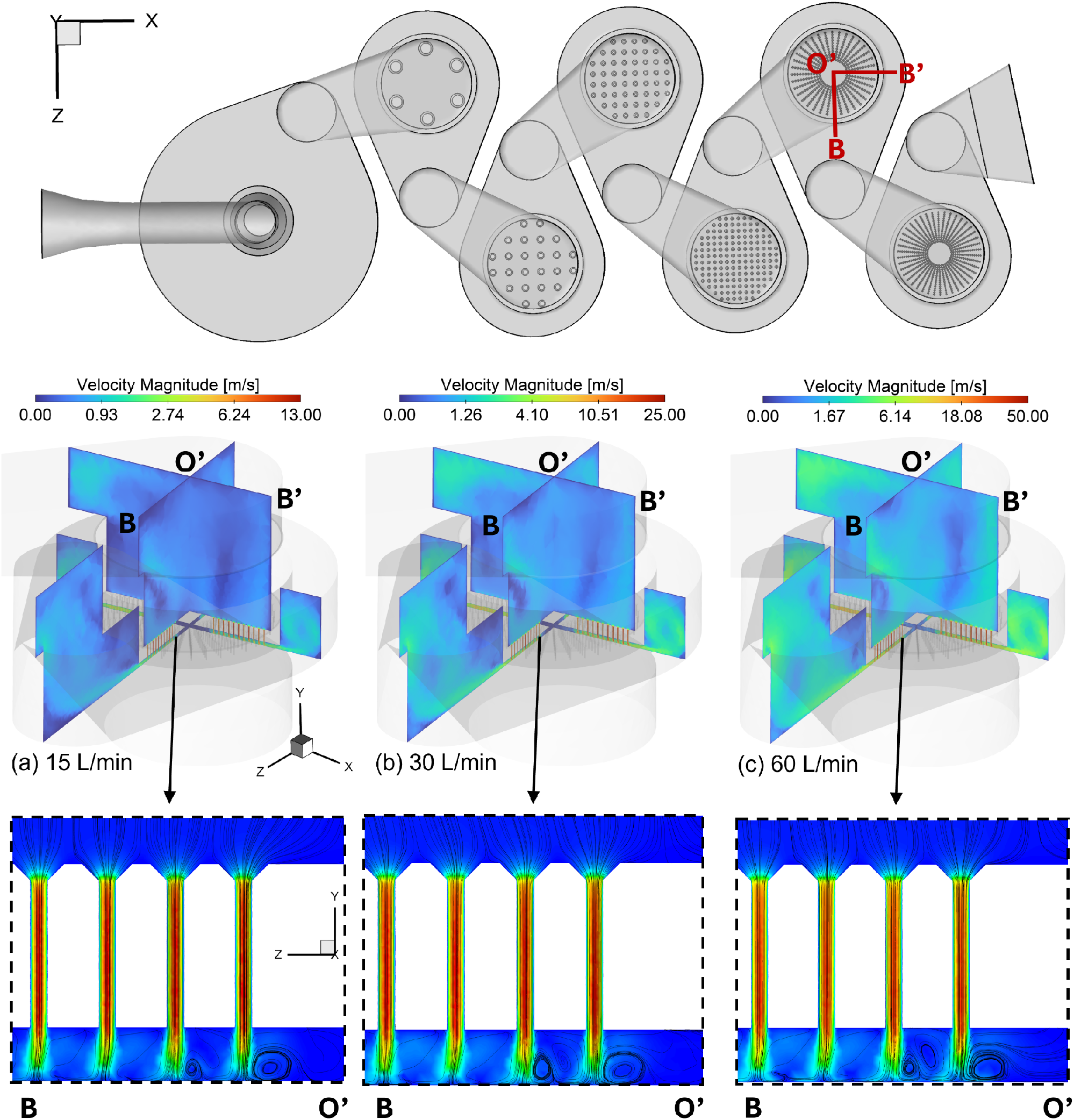
Velocity magnitude contours and streamlines at the filter of Stage 6 (S6) at three different flow rates: (a) 15 L/min, (b) 30 L/min, and (c) 60 L/min

**Figure 10.**
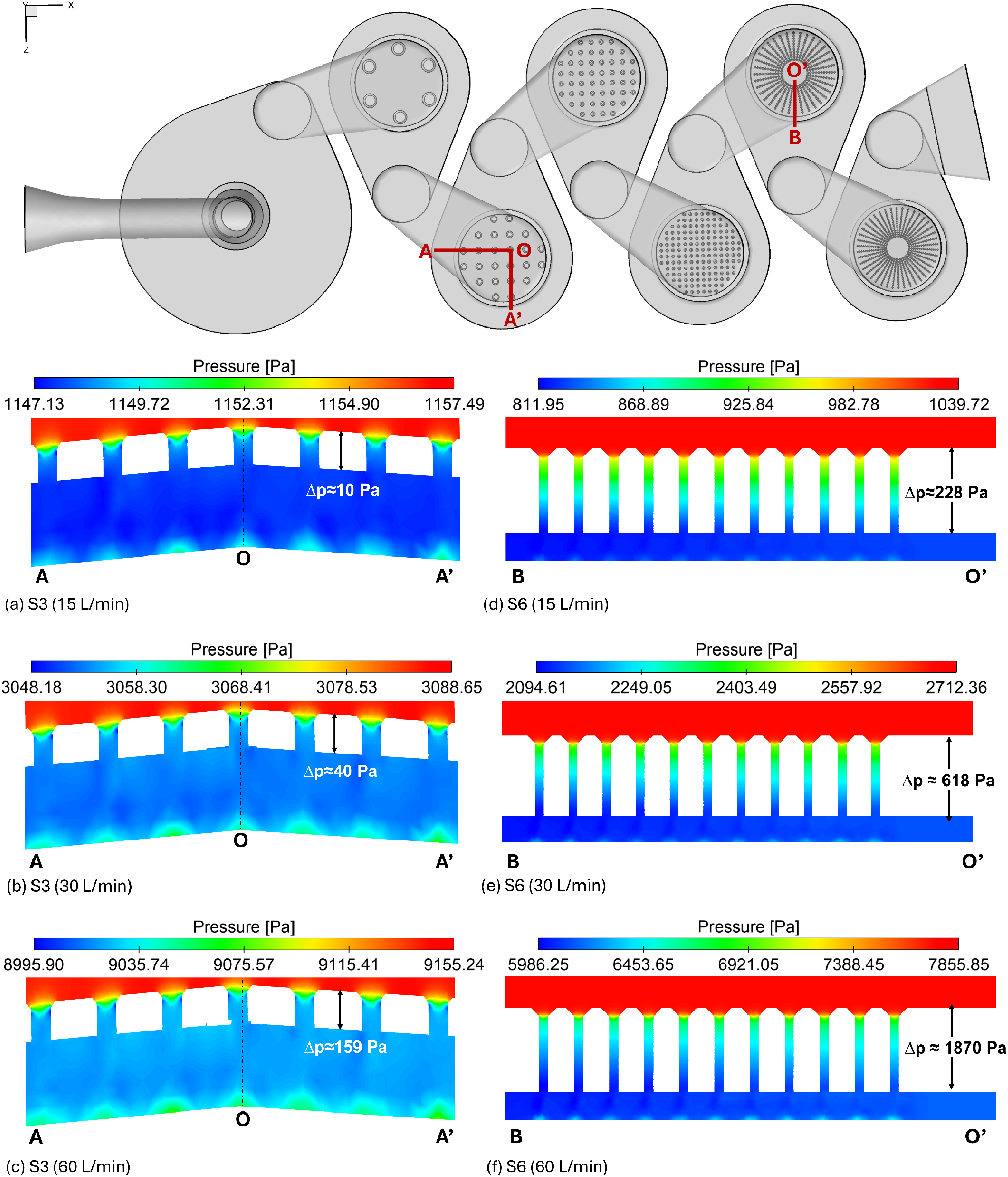
Pressure contours near the filters of S3 and S6 with three different flow rates: (a) S3 at 15 L/min, (b) S3 at 30 L/min, (c) S3 at 60 L/min, (d) S6 at 15 L/min, (e) S6 at 30 L/min, and (f) S6 at 60 L/min

The crossflow phenomenon observed in the multi-nozzle impactor studied by Fang et al. [35] can be effectively evaluated by analyzing the turbulent intensity (*TI*). It can be seen from **Fig. 11** that, for both S3 and S6, *TI* is not uniform across the nozzle filter and downstream at all flow rates. Specifically, *TI* is higher in the filter nozzles closer to the center of the stages than at the peripheral nozzles. For S3, the maximum *TI* between the nozzle filter and the impaction plate is about 0.2, 0.57, and 1.4 at 15, 30, and 60 L/min.

**Figure 11.**
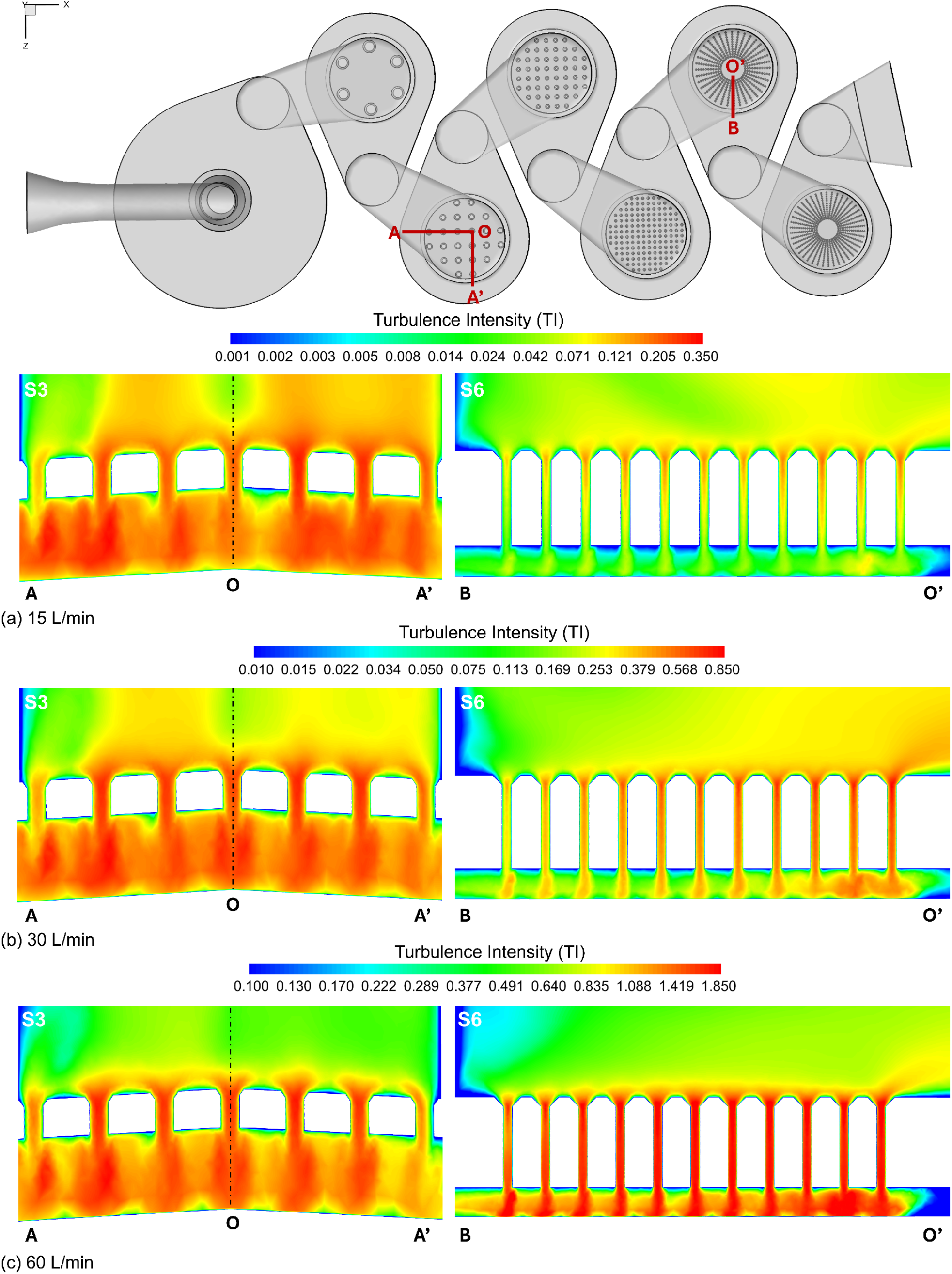
Turbulence intensity (*TI*) contours near the filters of S3 and S6 with three different flow rates: (a) 15 L/min, (b) 30 L/min, and (c) 60 L/min

At S6, the maximum *TI* decreased to 0.008, 0.5, and 1.4 at the respective flow rates. Comparatively, *TI* is higher at S3 than at S6 with all flow rates. Based on velocity contours (see **Figs. 8 and 9**) and *TI* (see **Fig. 11**), it can be found that although research findings [26, 60, 66] suggest that to keep *S*/*W* > 1 for round jet inertial impactors, this CFPD study shows that even *S*/*W* > 3 is not sufficient to entirely eliminate the cross flow phenomenon for S3 and S6 at all flow rates. Another indirect way to analyze the crossflow is to examine the impaction region in terms of wall shear stress (*WSS*). These impaction regions are the most likely surface locations for particle inertial impaction under the strong influence of unidirectional flow from the nozzle towards the impaction stage. Considering the jet-to-plate distance *S*, it can be roughly concluded that the wider the impaction regions, the higher the likelihood of crossflow, as the superposition of one nozzle’s impaction region with another nozzle’s neighboring impaction region indicates crossflow. For S6 and S7, such observations are seen at all flow rates (see **Fig. 12**), predominantly near the center of the impactor stage. The highest *W* and S3 are distinct and *SS* regions are more likely to interfere with the neighboring nozzle exits, and this likelihood increases with flow rate. One interesting observation is that as the flow rate increases, *WSS* increases as expected due to the increasing stagewise *Re*. The maximum *WSS* observed for both the stages for 15 L/min, 30 L/min, and 60 L/min is 5 Pa, 14 Pa, and 42 Pa respectively (see **Fig. 12**), S7 has *S*/*W* = 4.9 which is about 1.58 times higher than the S6, with both having same jet-to-plate distance (*S*) (see **Fig. 1(e)**) and marginally lowering the *Re* by 0.6%, 1.2%, and 1.5% when compared each other for S6 and S7 (see **Table 4**) with respective decreasing inhalation flow rates. This allowing the airstream to get enough room for the flow separation, increasing the interference in impaction regions on the impaction plate due to increased number of nozzles (*K*) in the sieve with same cluster diameter (*Dc*) of 38 *mm* (see **Fig. 1(f)**), contributing to 28 %, 31%, and 33% increase in average *WSS* for S7 when compared with S6 for 15 L/min, 30 L/min, and 60 L/min, respectively. Therefore, it can be concluded that only considering the *S*/*W* is not sufficient to avoid crossflow in the complex multi-nozzle impactor structure. The progressive nozzle pitch, higher at the center and lower towards the periphery of the nozzle cluster, could help mitigate crossflow effects, as crossflow intensity is greater at the impactor stage center than at the periphery.

**Figure 12.**
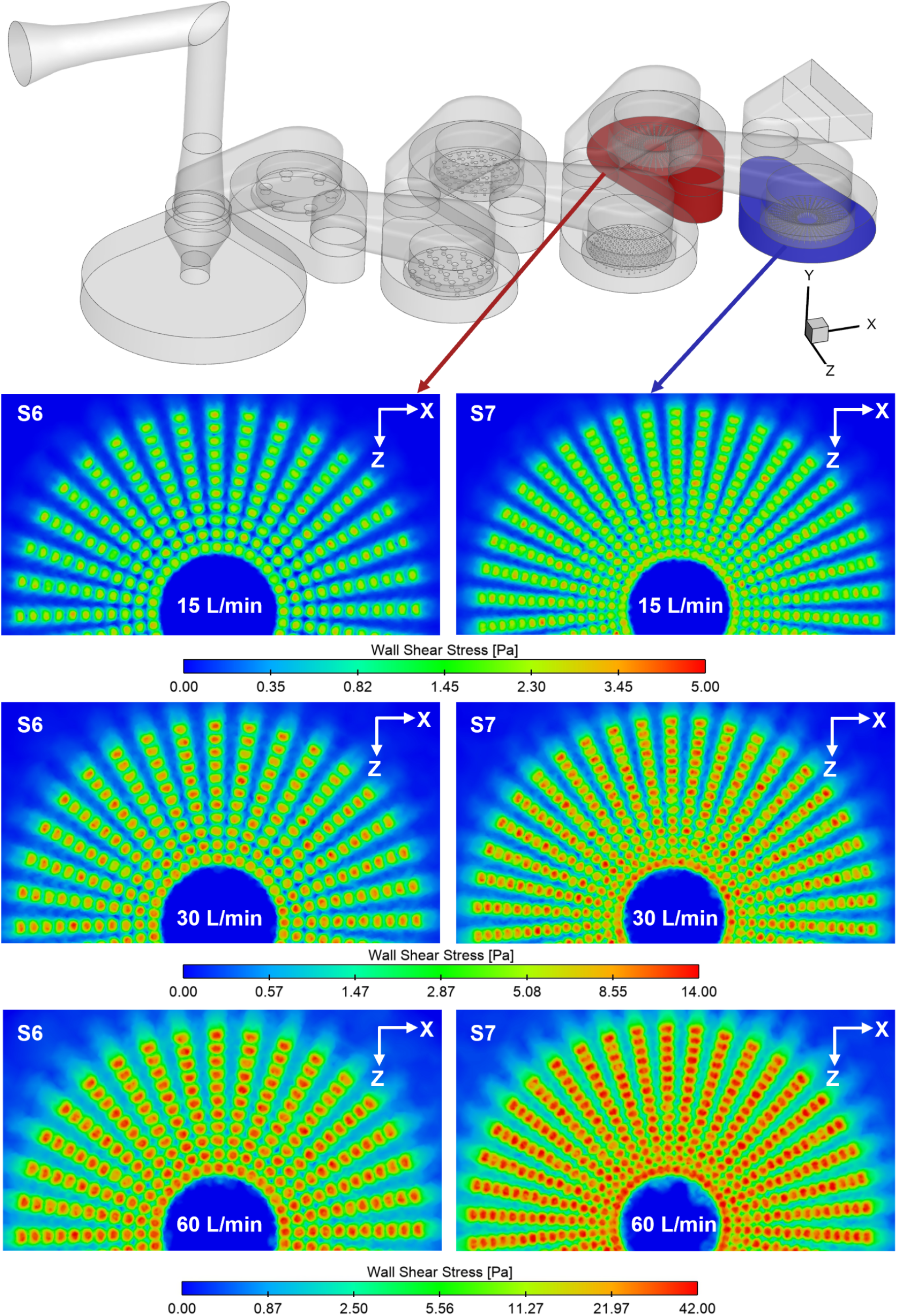
Bottom Wall shear stress (*WSS*) contours for Stage 6 (S6) on the left and Stage 7 (S7) on the right of the vNGI at three inhalation flow rates.

### 3.2 Monodisperse Particle Deposition and Interstage Wall Losses

To understand how particle size and flow rate influence particle deposition, collection efficiency, cutoff diameter, and wall losses, monodisperse particle simulations were first conducted at flow rates from 15 L/min to 60 L/min in vNGI with USP IP. The steady-state flow fields (see **Section 3.1**) were used for subsequent DPM simulations, in which 10,000 monodisperse particles were injected with aerodynamic diameters ranging from 0.2 to 20.0 µm. The collection efficiency (*CE*) and cutoff diameter (*d*_50_), were predicted for both flow rates. The *CEs* for 15, 30, and 60 L/min are shown in **Fig. 13 (b), Fig. 7(b)**, and **Fig. 13(d)**, respectively. Stagewise *d*_50_ at the increasing flow rate are shown in **Fig. 13 (a), Fig. 7 (a), and Fig. 13(c)**, respectively.

**Figure 13.**
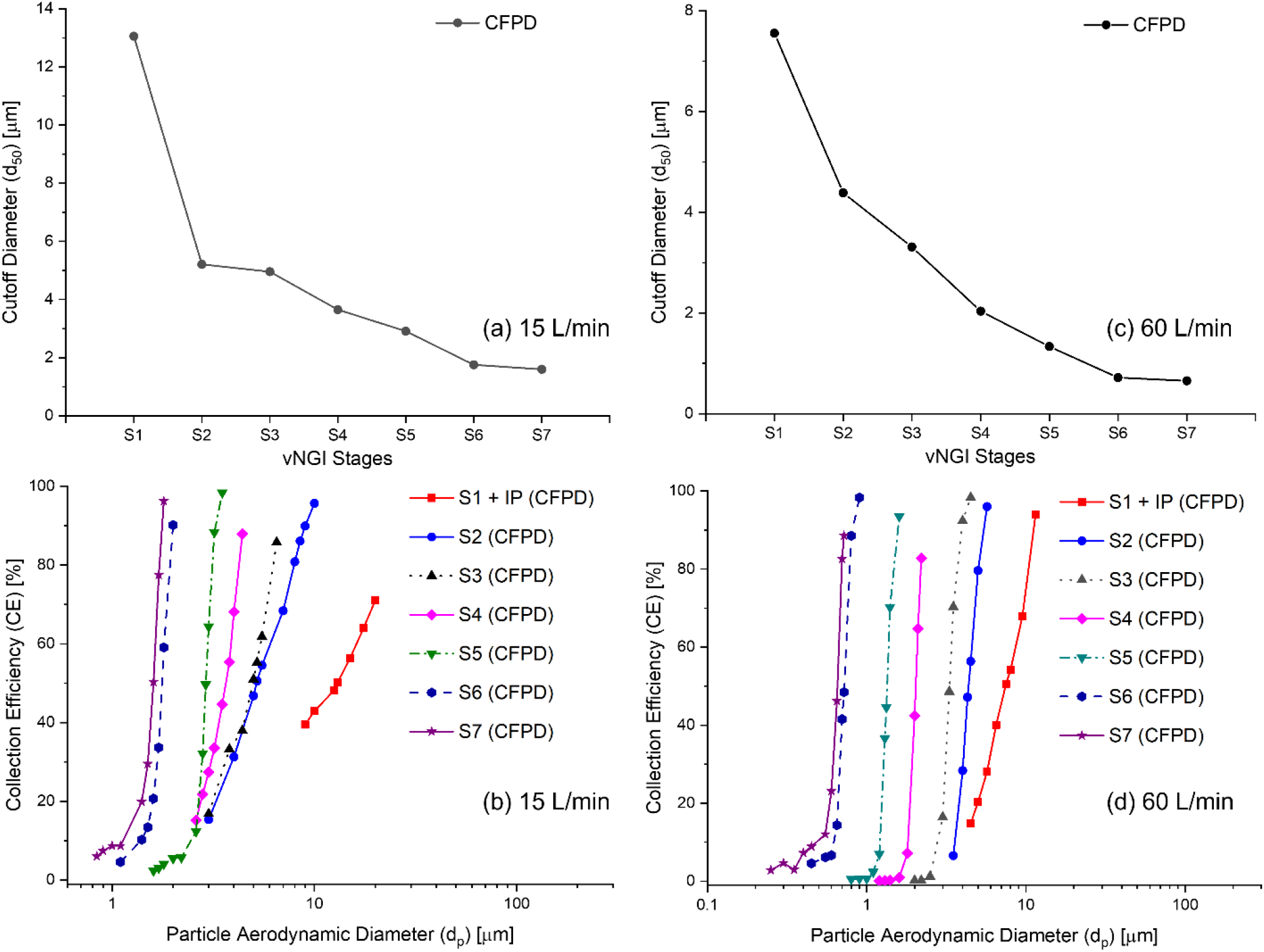
CFPD predicted cutoff diameters (*d*_50_) and particle collection efficiencies (*CE*) in vNGI stages with different flow rates: (a) Cutoff diameters at 15 L/min, (b) Collection efficiencies at 15 L/min, (c) Cutoff diameters at 60 L/min, and (d) Collection efficiencies at 60 L/min

For 15 L/min (see **Fig. 13(b)**), the *CE* curves for S1-S3 are widespread over the range of diameters, while the other stages have given a good, sharp cut-off. Similar to the 30 L/min (see **Fig. 7(b)**), the *CE* curves for S6 and S7 are almost superimposed on each other. For 60 L/min specifically, *CE* values for all the stages are distinct, i.e., no stagewise *CE*s overlap, which is also observed at the other two flow rates. The *CE* curve of S7 is seen nearly superimposed with *CE* curve of S6. All stages have shown sharper *CE* curves at 60 L/min than at the other two lower steady flow rates analyzed in this study. When *CEs* are analyzed along with the *d*_50_, the cut-off sharpness can be understood well. For example, at 15 L/min (see **Figs. 13(a)** and **(b)**) and 30 L/min (see **Fig. 7**), S2 and S3 have shown *d*_50_ nearly the same (i.e., 4 µm and 4.5 µm). Additionally, *CEs* for the same stages can be seen overlapping near to the 50% of *CE*. Based on the discussion in **Section 2.6.2**, the vNGI model has underperformed for S2 by about 25% in case of *d*_50_ than the experimental results [9] at 30 L/min. A similar phenomenon could have occurred at 15 L/min, shifting *CE* curve towards the left (see **Fig. 13(b)**). For 60 L/min (see **Figs. 13 (c) and (d)**), *d*_50_ at S2 and S3 are distinct and, *CEs* are also not overlapped, demonstrating a good example of sharp cutoff. It indicates that higher flow rates yield better cutoffs, especially for S1 and S2. The overlapping of *CEs* for 15 L/min and 30 L/min cases could be because of NGI design. S1 nozzle develops the highest *Re* and has highest area for particle impaction (see **Fig.1**), which is a suitable environment for developing recirculation zones, which can lead deposited particles to detach into the airflow called re-entrainment, more likely to occur on small diameter particles in reality. On the other hand, the vNGI considers particles that touch the surface to deposit, leaving no chance for them to re-entrain. In *CE* curves plotted for S1 and USP IP together would have captured the small diameter particles of stage S2, showing a sudden knee shape near 50% *CE* at the S1 + IP *CE* curve and interfering the *CE* of S2 at lower *CE* level, followed in all the flow rates for example in deposition of particle diameter of 5.5 µm for 30 L/min flow rate, the S1 + IP deposition is found to be 37% where the S2 deposition is 62.8% out of total deposited particles. At 15 L/min, it is 32.7% in S1+IP and 54.5%, and at 60 L/min, it is nearly 28% and 96% for S1+IP and S2 stages, respectively. Along with the increasing flow rate, the overlapping is decreased and so *d*_50_ as it can occur because the smaller particle tends to travel along with the airflow field and to keep the 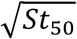 near to the 0.495 by NGI design [26], the particle diameter has to decrease when the exit velocity is increased as per the Eq. (46) and Eq. (47). Other similar studies with the Anderson cascade impactor have also shown the similar sudden knee shape for first two stages as S0 and S1 in Aderson cascade terminology, where USP IP fits at S0 and the consecutive stage is S1 [21, 22] for 28.3 L/min steady flow rate but the sudden change in *CE* is not discussed because it may not be that distinct than the current study.

Particle deposition on areas other than the collection cup stages (see **Fig. 1(e)**) and the USP IP (or other MT models shown in **Fig. 3**) can be defined as the interstage wall loss (*WL*) [69, 70]. Although minimal *WL* in NGI is reported [26, 71, 72] with respect to the total dose injected at the inlet, this study found a significant fraction of *WL* is due to the concentrated deposition near the jet entry, i.e., the peripheral area of the nozzle where the upstream flow enters the nozzle and bends, which can be seen in insets of **Figs. 19 (a)-(d)**. Similar observations have been noted in previous studies of the Andersen cascade impactor [21, 70]. Monodisperse particle *WL* at different flow rates for each stage are shown in **Figs. 14 (a)-(g)**. It does not include MOC, since it is modeled only partially in this study. The particle wall loss shown in **Fig. 14** can be considered as interstage *WL* at the particular stage when compared with the previous stage. For example, **Fig. 14(b)** represents the *WL* between the S1 and S2 for analysis of *WL* for S2. The interstage *WL* can be calculated as

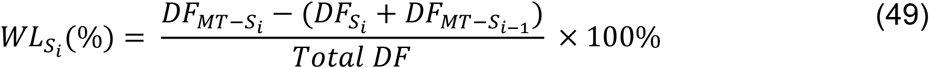

where *S*_*i*_ and *S*_*i*−1_ are the consecutive stages.

**Figure 14.**
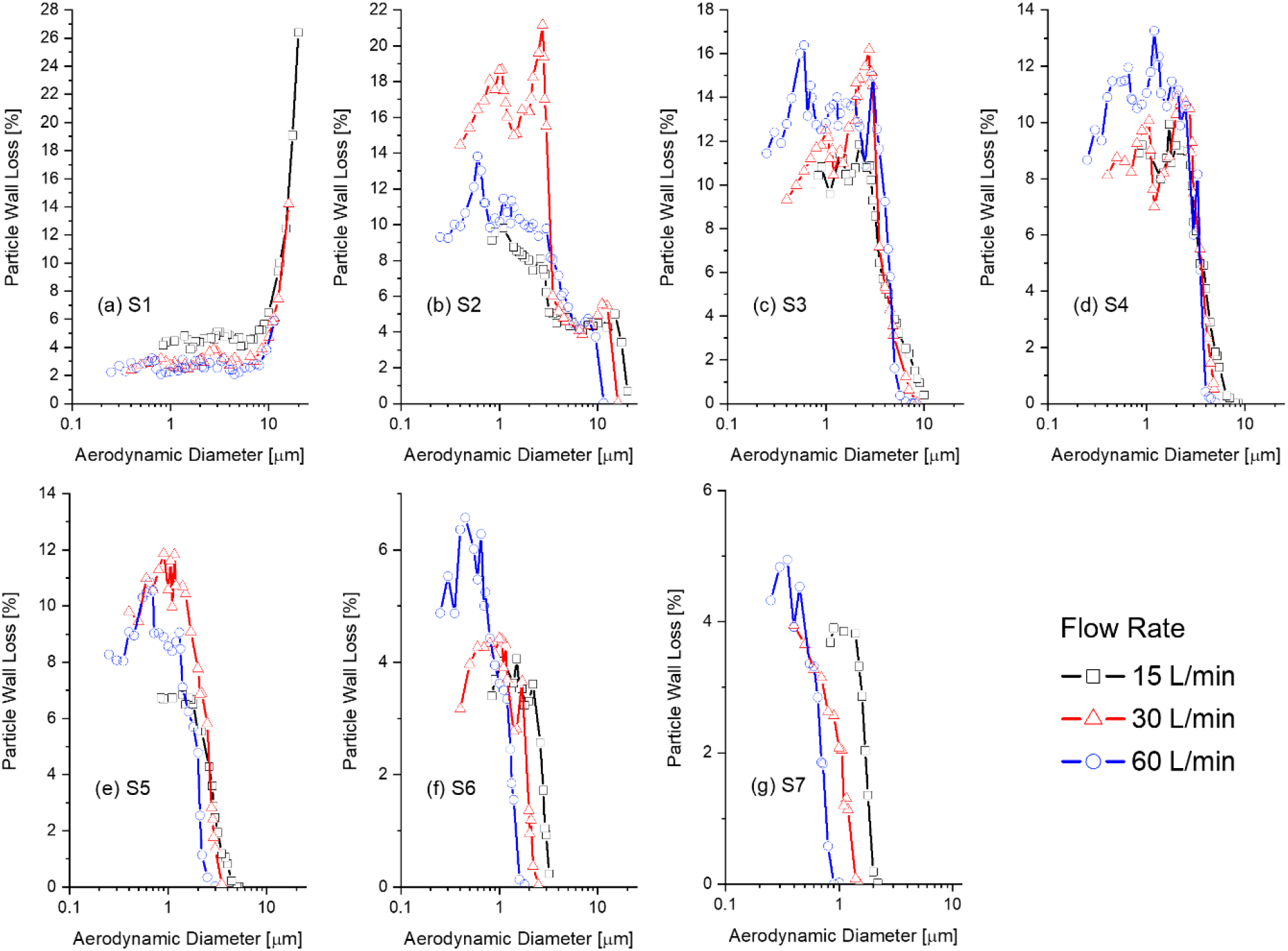
Particle wall losses (%) at steady-state flow rates of 15, 30, and 60 L/min across vNGI stages: (a) S1 (b) S2, (c) S3, (d) S4, (e) S5, (f) S6, and (g) S7

Overall, the maximum interstage *WL* ranges from 5% to 26%, depending on the flow rates and stage of consideration. On the other hand, if the cumulative *WL* for the particles with *d*_*ae*_ <1 µm across all the stages is analyzed, the overall cumulative *WL* reaches 45-49%, 53-62%, and 50-64%, as the flow rate increases. This could be due to intensified turbulence dispersion at higher flow rates, along with the assumption of a perfect-trap wall boundary condition for the particle that touches the wall without rebound.

At S1, the major *WL* is seen at the converging section from USP IP to the S1 filter nozzle for large particles (see **Fig. 14(a)**), as it is the only region where the deposition of particles is not considered as stage deposition of USP IP or S1. For particles smaller than 7 µm, the *WL* is below 5% for 15 L/min. As the flow rate increases, *WL* decreases. It can be observed that higher flow rates result in lower *WL* than at low flow rates due to the nozzle’s converging area, which creates a favorable pressure gradient that promotes streamlined flow by reducing flow separation. Additionally, as particle size increases beyond 7 µm, the overall trend of *WL* shows a rapid increase due to gravitational influence and the particle’s higher momentum, leaving insufficient time for the particle to move along the streamlines in the converging section.

From S2 and onwards, the trend of *WL* vs. particle size shifts in the opposite direction compared with S1 due to cut-offs of S1, where large diameters near 10 µm are filtered out based on operating flow rate conditions. The *WL* from S2 onwards is largely contributed by the upper part of the nozzle and a connecting flow channel which connects collection cup of a stage to the nozzles of the next stage with a converging to diverging section (see **Figs. 1 (a), (b), and (e)**) creating unfavorable pressure gradient, reducing flow field velocity, and giving rise to flow separation for a particle laden flow. In addition, the flow field suddenly passes through several small nozzles. Due to these structural design features of NGI, a case of backward-facing step is developed in which the particle-laden flow experiences turbulent dispersion and resultant *WL*.

In S2 at 30 L/min, *WL* is observed to be the highest among all flow rates (see **Fig. 14(b)**). This anomalous behavior for particles near and lower than 1 µm could have had a major effect on the *CEs* for S4 onwards (see **Fig. 7(b)**). With the increase in stage numbers for all flow rates, *WL* decreases (see **Fig. 14 (c)-(g)**) as the number of filter nozzles increases, thereby supporting flow regularization. For similar reasons, overall, stage-wise *WL* decreases as the flow rate decreases, except for S1. This is due to turbulence dispersion and inertial impaction-induced deposition at lower flow rates in the interstage sections. The S1 filter nozzle is a type of single round nozzle achieving the highest *Re* compared to others, which is susceptible to collecting larger diameter particles (> 7 µm) due to inertial impaction rather than diffusion for smaller diameter particles.

The discussion in this section on monodisperse particle deposition demonstrates the NGI’s design attribute based on theoretical considerations to filter out larger particles at the start of the stage. The exact quantification of *WL* with respect to stages and flow rate was not found. However, the general trend of interstage *WL* can be seen from **Fig. 14**. The analysis on interstage *WL* shows that smaller-diameter particles can deposit before their designated sampling stage, and this can similarly propagate into APSD measurements, largely reducing particle deposition and having lower *MAD*/*CMD*, which is discussed in the next section. It is worth mentioning that *WL* is a major challenge for impactor designers [68] in the assessment of APSD, which could be addressed by using the vNGI presented in this study.

### 3.3 Polydisperse Particle Deposition

Polydisperse aerosols contain a range of particle diameters that can be characterized by *MAD* or *CMD*, with *GSD* describing the distribution spread. In this study, four *CMD* − *GSD* combinations (see **Fig. 4** and **Table 3**) were employed to predict particle deposition fraction (*DF*) in vNGI at 15, 30, and 60 L/min (see **Figs. 15 (a)-(c)**). In this study, the particles equal to and below 1.0 µm were modeled additionally with the Brownian motion (see **Section 2.2.2**), and particles exiting from the vNGI through the outlet (see **Fig. 1**) were assumed to be deposited in the MOC, as by design it is collecting all remaining undeposited particles downstream to S7. **Table 5** contains the sum of depositions in four categories for all the flow rates, i.e., (1) USP IP-to-S2, (2) S3-to-S5, (3) S6-to-MOC regions, and (4) the total deposition across stages. The idea of grouping the stagewise deposition is inspired by previous studies [68, 73, 74], and the simplicity of analysis.

**Table 5.**
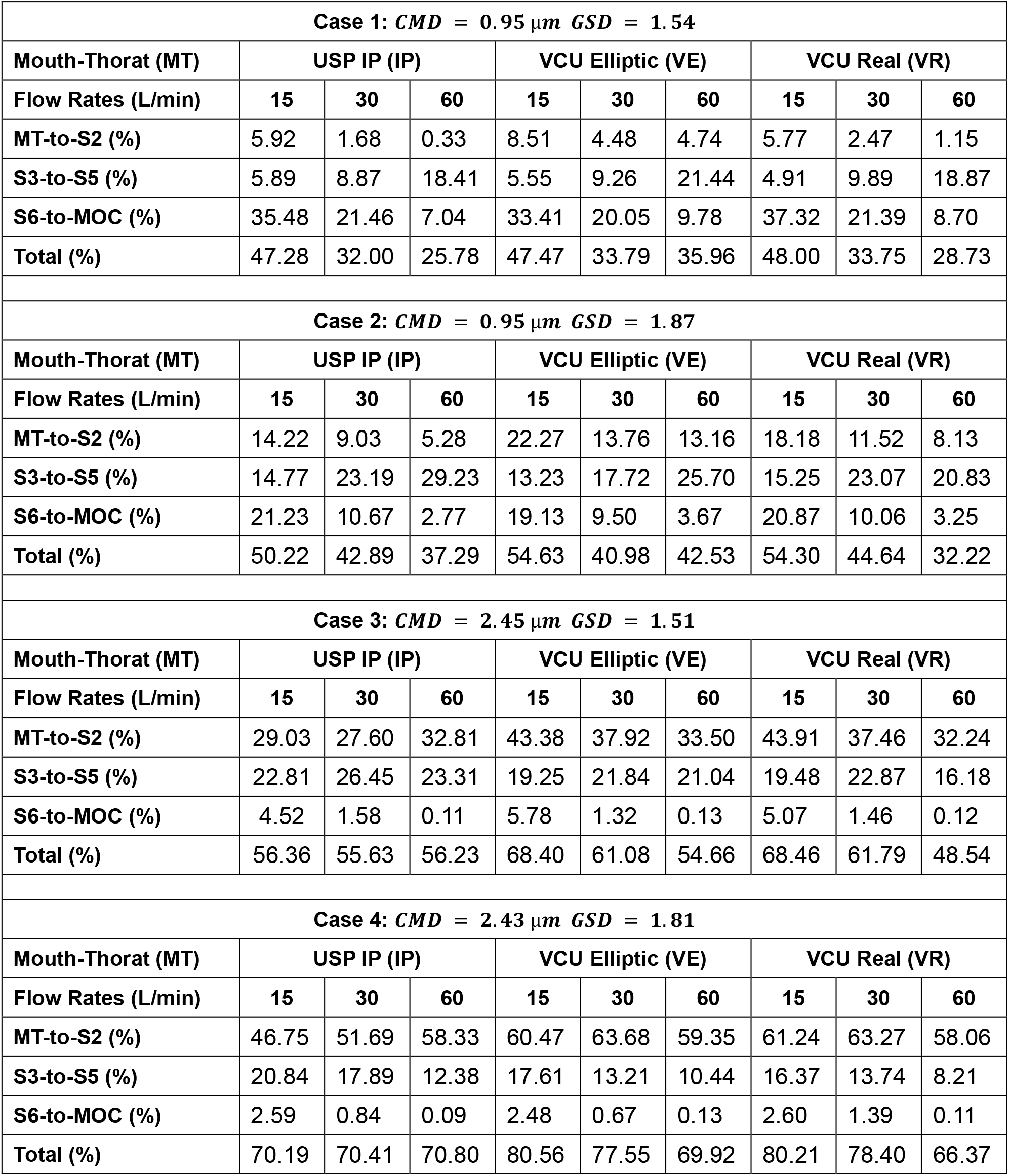
CFPD predicted particle deposition fraction (*DF*) across the different MT models at all the flow rates.

| Case 1: $CMD = 0.95 \mu m$ $GSD = 1.54$ | | | | | | | | | |
| --- | --- | --- | --- | --- | --- | --- | --- | --- | --- |
| Mouth-Thorax (MT) | USP IP (IP) |  |  | VCU Elliptic (VE) |  |  | VCU Real (VR) |  |  |
| Flow Rates (L/min) | 15 | 30 | 60 | 15 | 30 | 60 | 15 | 30 | 60 |
| MT-to-S2 (%) | 5.92 | 1.68 | 0.33 | 8.51 | 4.48 | 4.74 | 5.77 | 2.47 | 1.15 |
| S3-to-S5 (%) | 5.89 | 8.87 | 18.41 | 5.55 | 9.26 | 21.44 | 4.91 | 9.89 | 18.87 |
| S6-to-MOC (%) | 35.48 | 21.46 | 7.04 | 33.41 | 20.05 | 9.78 | 37.32 | 21.39 | 8.70 |
| Total (%) | 47.28 | 32.00 | 25.78 | 47.47 | 33.79 | 35.96 | 48.00 | 33.75 | 28.73 |
| Case 2: $CMD = 0.95 \mu m$ $GSD = 1.87$ | | | | | | | | | |
| Mouth-Thorax (MT) | USP IP (IP) |  |  | VCU Elliptic (VE) |  |  | VCU Real (VR) |  |  |
| Flow Rates (L/min) | 15 | 30 | 60 | 15 | 30 | 60 | 15 | 30 | 60 |
| MT-to-S2 (%) | 14.22 | 9.03 | 5.28 | 22.27 | 13.76 | 13.16 | 18.18 | 11.52 | 8.13 |
| S3-to-S5 (%) | 14.77 | 23.19 | 29.23 | 13.23 | 17.72 | 25.70 | 15.25 | 23.07 | 20.83 |
| S6-to-MOC (%) | 21.23 | 10.67 | 2.77 | 19.13 | 9.50 | 3.67 | 20.87 | 10.06 | 3.25 |
| Total (%) | 50.22 | 42.89 | 37.29 | 54.63 | 40.98 | 42.53 | 54.30 | 44.64 | 32.22 |
| Case 3: $CMD = 2.45 \mu m$ $GSD = 1.51$ | | | | | | | | | |
| Mouth-Thorax (MT) | USP IP (IP) |  |  | VCU Elliptic (VE) |  |  | VCU Real (VR) |  |  |
| Flow Rates (L/min) | 15 | 30 | 60 | 15 | 30 | 60 | 15 | 30 | 60 |
| MT-to-S2 (%) | 29.03 | 27.60 | 32.81 | 43.38 | 37.92 | 33.50 | 43.91 | 37.46 | 32.24 |
| S3-to-S5 (%) | 22.81 | 26.45 | 23.31 | 19.25 | 21.84 | 21.04 | 19.48 | 22.87 | 16.18 |
| S6-to-MOC (%) | 4.52 | 1.58 | 0.11 | 5.78 | 1.32 | 0.13 | 5.07 | 1.46 | 0.12 |
| Total (%) | 56.36 | 55.63 | 56.23 | 68.40 | 61.08 | 54.66 | 68.46 | 61.79 | 48.54 |
| Case 4: $CMD = 2.43 \mu m$ $GSD = 1.81$ | | | | | | | | | |
| Mouth-Thorax (MT) | USP IP (IP) |  |  | VCU Elliptic (VE) |  |  | VCU Real (VR) |  |  |
| Flow Rates (L/min) | 15 | 30 | 60 | 15 | 30 | 60 | 15 | 30 | 60 |
| MT-to-S2 (%) | 46.75 | 51.69 | 58.33 | 60.47 | 63.68 | 59.35 | 61.24 | 63.27 | 58.06 |
| S3-to-S5 (%) | 20.84 | 17.89 | 12.38 | 17.61 | 13.21 | 10.44 | 16.37 | 13.74 | 8.21 |
| S6-to-MOC (%) | 2.59 | 0.84 | 0.09 | 2.48 | 0.67 | 0.13 | 2.60 | 1.39 | 0.11 |
| Total (%) | 70.19 | 70.41 | 70.80 | 80.56 | 77.55 | 69.92 | 80.21 | 78.40 | 66.37 |

**Figure 15.**
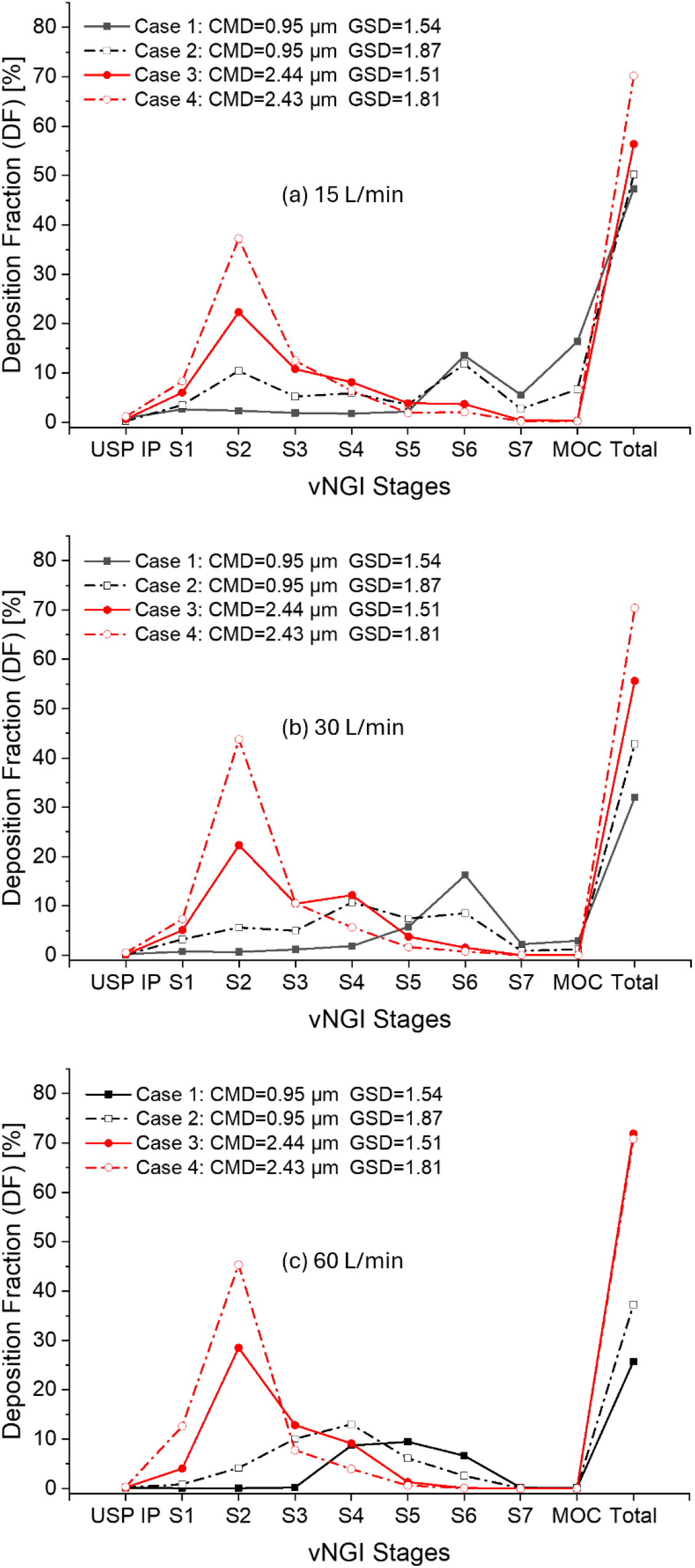
Regional deposition fractions (*DFs*) with USP IP mouth-throat model and vNGI stages at different flow rates: (a) 15 L/min, (b) 30 L/min, and (c) 60 L/min

Comparison of the stagewise *d*_50_ and *DF* values indicate that the vNGI model generally captured the deposition behavior of polydisperse aerosol-laden flows for cases in which more than 90% of the particles were smaller than 5 µm by number. However, interstage *WL*, like that observed for monodisperse aerosols in **Section 3.2**, remained an important factor affecting the total *DF*. As shown in **Table 5**, the total *DF* increased progressively from Case 1 to Case 4 at all flow rates, suggesting that interstage *WL* could have decreased as the particle size distribution shifted toward larger particle diameters.

As discussed earlier, interstage *WL* was more pronounced for particles smaller than 1.0 µm. Consequently, Cases 1 and 2, which contained a larger fraction of submicron particles, exhibited substantially lower total *DFs*. In contrast, Cases 3 and 4 could have been less affected by interstage *WL*, because their particle size distributions shifted toward larger diameters. Flow rate also influenced the particle deposition. At 15 L/min, particles had a longer residence time and were therefore better able to adjust to the local flow field, allowing greater penetration into the downstream NGI stages. This resulted in higher total *DFs* for Cases 1 and 2 than at higher flow rates. As the flow rate increased, the total *DF* generally decreased.

The increasing trend in *DF* from Case 1 to Case 4 can be attributed to the progressive increase in particle size and the broadening of particle size distribution. Cases 1 and 2 corresponded to distributions with *CMD* ≈ 0.95 μ*m* but different mass median diameters, *MMD* ≈ 1.6 μ*m* and 2.6 μ*m*, respectively. Cases 3 and 4 represented larger particle distributions with *CMD* ≈ 2.5 μ*m* and *MMD* ≈ 4.0 μ*m* and 5.6 μ*m*, respectively, with increasing *GSD* (see **Table 3**). Therefore, the case-dependent increase in total *DF* was primarily associated with reduced interstage *WL* and enhanced stage deposition as the particle size distribution shifted toward larger diameters.

Cases 1 and 2 were more sensitive to increasing flow rate, as reflected by the reduction in *DF* in the USP IP-to-S2 and S6-to-MOC regions (see **Table 5**). This trend was particularly evident at the MOC (see **Fig. 15**). In contrast, the intermediate stages (i.e., S3-to-S5 region) generally showed an increasing *DF* with increasing flow rate. Compared with Case 1, which had an *MMD* of approximately 1.6 μ*m*, Case 2 had a larger *MMD* of approximately 2.6 μ*m* and a broader particle size distribution. Therefore, based on the stagewise *d*_50_ values, a larger fraction of particles in Case 2 was expected to deposit in the S3-to-S5 region, while fewer particles were expected to penetrate to the S6-to-MOC region (see **Figs. 7 (a), 13 (a), and 13 (c)**). This behavior is consistent with the combined cut-off range of S3 to S5, which shifted from 4.95-2.90 µm at 15 L/min to 4.21-2.02 µm at 30 L/min and 3.31-1.34 µm at 60 L/min. Similarly, the *d*_50_ of S6 decreased from 1.76 µm to 1.17 µm and 0.70 µm as the flow rate increased, allowing this stage to collect the remaining smaller particles.

For Cases 3 and 4, the *DF* in the USP IP-to-S2 region increased with flow rate, whereas the *DF* in the S6-to-MOC region decreased (see **Table 5**). The behavior of the intermediate stages, i.e., S3-to-S5 region, was more case-dependent. In Case 3, the *DF* in S3-to-S5 region varied only marginally among the three flow rates, with changes ranging from 0.5% to 3.64%. However, in Case 4, the *DF* in S3-to-S5 region decreased with increasing flow rate. Case 3 had an *MMD* of approximately 4.0 µm, whereas Case 4 had an *MMD* of approximately 5.7 µm. The corresponding *GSD* were similar, approximately 1.5 and 1.6, respectively. Based on the stagewise *d*_50_, most particles in Cases 3 and 4 were expected to deposit in S2, whose cut-off diameter decreased from 5.19 µm at 15 L/min to 4.89 µm at 30 L/min and 4.36 µm at 60 L/min (see **Figs. 7 (a), 13 (a)**, and **13 (c)**). This expected deposition pattern was well captured by the vNGI, as shown in **Fig. 15** and **Table 5**.

In general, the *DFs* in the USP IP-to-S1 region were low because the *d*_50_ values of these upstream stages were larger than approximately 4 µm, which is close to the cut-off diameter of S2 (see **Figs. 7 (a), 13 (a), and 13 (c)**). Therefore, only particles much larger than 4 µm were likely to deposit in the USP IP-to-S1 region, while smaller particles were transported to the downstream stages at all flow rates. Cases 1 and 2 contained only a small fraction of particles larger than or equal to 4 µm by number, with values of less than 1% and 0.64%, respectively. However, these particles accounted for 0.64% and 23.77% of the total aerosol mass, respectively. In contrast, Cases 3 and 4 contained 13.61% and 21.75% of particles larger than or equal to 4 µm by number, corresponding to 52.76% and 77.66% of the total aerosol mass, respectively.

For Cases 1 and 2, the predicted *DF* values in the USP IP-to-S1 region were less than or equal to 1% at 30 and 60 L/min. Comparatively, at 15 L/min, the *DF* increased to approximately 3.6%–3.8% as the particle size distribution broadened from Case 1 to Case 2. Within this upstream region, the USP IP alone collected 0.90% and 0.27% of the aerosol for Cases 1 and 2, respectively (see **Fig. 15** and **Table 6**). For Cases 3 and 4, the *DF* values in the USP IP-to-S1 regions generally decreased from low to high flow rate, with values of 6.7% and 9.5% at 15 L/min, 5.3% and 7.9% at 30 L/min, and 4.3% and 13.0% at 60 L/min for Cases 3 and 4, respectively. The exception was Case 4 at 60 L/min, where S1 alone accounted for 12.63% in *DF*, while the USP IP contributed only a marginal fraction.

**Table 6.** CFPD predicted particle deposition fraction (*DF*) across the different MT configurations at all the flow rates in MT and S1.

| Case 1: $CMD = 0.95 \mu m$ $GSD = 1.54$ | | | | | | | | | |
| --- | --- | --- | --- | --- | --- | --- | --- | --- | --- |
| Mouth-Thorat (MT) | USP IP (IP) |  |  | VCU Elliptic (VE) |  |  | VCU Real (VR) |  |  |
| Flow Rates (L/min) | 15 | 30 | 60 | 15 | 30 | 60 | 15 | 30 | 60 |
| MT (%) | 0.90 | 0.25 | 0.19 | 3.43 | 3.06 | 4.17 | 0.01 | 1.12 | 0.95 |
| S1 (%) | 2.66 | 0.74 | 0.07 | 3.44 | 0.91 | 0.00 | 2.68 | 0.62 | 0.00 |
| S2 (%) | 2.37 | 0.68 | 0.07 | 1.64 | 0.51 | 0.56 | 3.08 | 0.73 | 0.20 |
| Case 2: $CMD = 0.95 \mu m$ $GSD = 1.87$ | | | | | | | | | |
| Mouth-Thorat (MT) | USP IP (IP) |  |  | VCU Elliptic (VE) |  |  | VCU Real (VR) |  |  |
| Flow Rates (L/min) | 15 | 30 | 60 | 15 | 30 | 60 | 15 | 30 | 60 |
| MT (%) | 0.27 | 0.11 | 0.23 | 3.42 | 3.48 | 4.56 | 1.49 | 1.58 | 1.02 |
| S1 (%) | 3.52 | 3.27 | 0.87 | 13.17 | 7.82 | 1.44 | 9.07 | 7.04 | 0.63 |
| S2 (%) | 10.42 | 5.65 | 4.19 | 5.68 | 2.47 | 7.16 | 7.62 | 2.90 | 6.49 |
| Case 3: $CMD = 2.45 \mu m$ $GSD = 1.51$ | | | | | | | | | |
| Mouth-Thorat (MT) | USP IP (IP) |  |  | VCU Elliptic (VE) |  |  | VCU Real (VR) |  |  |
| Flow Rates (L/min) | 15 | 30 | 60 | 15 | 30 | 60 | 15 | 30 | 60 |
| MT (%) | 0.68 | 0.18 | 0.23 | 4.08 | 4.08 | 5.76 | 2.08 | 1.61 | 3.58 |
| S1 (%) | 6.03 | 5.08 | 4.06 | 21.55 | 19.90 | 6.08 | 22.93 | 21.71 | 7.39 |
| S2 (%) | 22.32 | 22.34 | 28.52 | 17.75 | 13.94 | 21.66 | 18.90 | 14.15 | 21.27 |
| Case 4: $CMD = 2.43 \mu m$ $GSD = 1.81$ | | | | | | | | | |
| Mouth-Thorat (MT) | USP IP (IP) |  |  | VCU Elliptic (VE) |  |  | VCU Real (VR) |  |  |
| Flow Rates (L/min) | 15 | 30 | 60 | 15 | 30 | 60 | 15 | 30 | 60 |
| MT (%) | 1.18 | 0.56 | 0.36 | 4.30 | 5.54 | 13.54 | 2.69 | 4.07 | 8.50 |
| S1 (%) | 8.33 | 7.35 | 12.63 | 25.08 | 28.72 | 20.10 | 28.64 | 34.84 | 25.37 |
| S2 (%) | 37.24 | 43.78 | 45.34 | 31.09 | 29.42 | 25.72 | 29.91 | 24.36 | 24.19 |

### 3.4 Mouth-Throat (MT) Variability Effect on vNGI Measurement

To investigate the MT configuration influence on vNGI measurement, three MT geometries shown in **Fig. 2** (i.e., USP IP, VCU Elliptic, and VCU Realistic) were employed for simulations using the four polydisperse particle size distributions (see **Fig. 4** and **Table 3**) at all three flow rates. Comparisons of *DFs* are shown in **Figs. 16 (a)-(d)**, and **Tables 5-6**.

**Figure 16.**
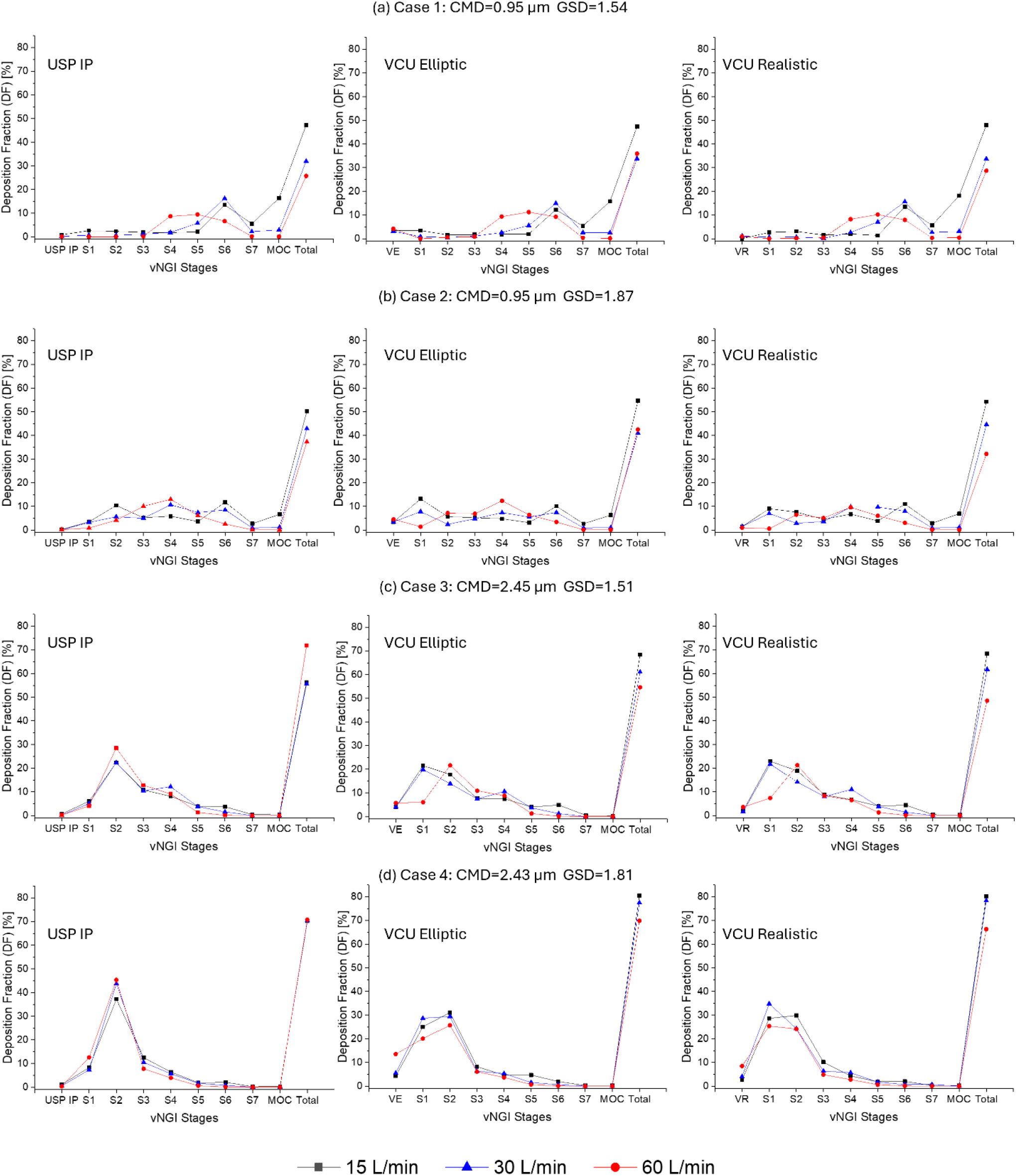
Regional deposition fractions (*DFs*) in vNGI with different mouth-throat geometries (i.e., USP IP (left), VCU Elliptic (VE, middle), VCU Real (VR, right)) with four polydisperse particle size distributions: (a) Case 1, (b) Case 2, (c) Case 3, and (d) Case 4.

First, it can be observed that 19 out of 24 configurations of bio-relevant MT geometries (i.e., VCU Elliptic and Realistic) have shown higher total *DF* than using USP IP (see **Fig. 16** and **Table 5**). All the lower total *DF* is observed for 60 L/min flow rate and with Case 3-4 for both bio-relevant MT and Case 2 for only VCU Realistic MT. As compared with total DF of USP IP MT, VCU Realistic MT showed the lowest total *DF* about 5%, 7.7%, and 4.4% for Cases 2-4, respectively. The VCU Elliptic MT showed 1.6% and 0.9% less total *DF* compared to USP IP MT for Cases 3 and 4 respectively. The discrepancy of lower total *DF* for VCU Real than VCU Elliptic could be because of suspending particles which are more prominent than the USP IP. As mentioned earlier, *WL* is the interstage wall deposition other than the required stage locations (see **Section 3.2**). Additionally, the remining amount from the total *DF* is contributed by the interstage WL and the suspended particles. In this study, interstage *WL* is not studied for polydisperse aerosol, as the underlying mechanism for *WL* such as excessive turbulence dispersion and accretion of particles at the peripheral entry region of the nozzle (see **Figs. 1(b)** and **(e)**) is assumed to remain consistent in the case of polydisperse aerosol.

Similar to the performance using USP IP, bio-relevant MT models also show an increasing trend of total *DF* with an increase in flow rate from Case 1 to 4 (see **Table 5**). Cases 1 and 2 show an increasing trend of *DF* for the S3-to-S5 region with respect to increasing flow rates, except for Case 2 for the 60 L/min flow rate, which showed *DF* lower than the 30 L/min flow rate by 2.2% but higher than the 15 L/min flow rate by 5.6% for the case of VCU Real MT. Furthermore, *DF* values for MT-to-S2 and S6-to-MOC regions show decreasing trend with respect to increasing flow rates. For Cases 3 and 4, the total *DF* for bio-relevant MT models decreases with increasing flow rates, a different behavior compared to vNGI with USP IP, where the total *DF* is almost unaffected. Also, *DF* values at bio-relevant MT-to-S2 and S6-to-MOC regions in Cases 3 and 4 is decreasing along with the increase in flow rate, except for both bio-relevant MT in Case 4 at 30 L/min, where it has increased slightly by 2-3% than 15 L/min flow rate for MT-to-S2 region. The *DF* for S3-to-S5 region in Case 3 showed a mixed response for VCU Real MT with a slight increment of 3.4% and decrement of 6.7% when compared between 15-to-30 L/min and 30-to-60 L/min respectively. The VCU Elliptic MT showed almost similar *DF* about 20.7% in average at S3-to-S5 region, less affected by the change in flow rates. With Case 4, both bio-relevant MT showed decreasing trend with increasing flow rates for *DF* at S3-to-S5 region. As shown in **Figs. 16 (a), 16 (b), and 17**, Cases 1 and 2 exhibited similar stagewise deposition trends between the bio-relevant MT models and the vNGI with the USP IP. In these two cases, most of *DF* accumulated in S5-to-S7 region at 15 and 30 L/min and in S4-to-S6 region at 60 L/min. In contrast, Cases 3 and 4 showed more noticeable differences among the MT models, particularly in the upstream stages S1 and S2 (see **Figs. 16 (c), 16 (d), and 17**). As discussed in **Section 3.3**, Cases 1 and 2 represent particle size distributions with smaller characteristic diameters, whereas Cases 3 and 4 represent distributions shifted toward larger particle diameters (see **Fig. 4** and **Table 3**). Accordingly, particles in Cases 1 and 2 were expected to deposit predominantly in the downstream stages of the vNGI, while particles in Cases 3 and 4 were expected to deposit preferentially in the MT and early stages based on cut-off diameters with respective flow rates. These results indicate that the bio-relevant MT geometries can substantially influence stagewise deposition in the vNGI, particularly for larger particle size distributions.

For further interpretation, **Figs. 16 and 17** and **Table 6** are considered together to compare the qualitative deposition patterns and quantitative *DFs* in the MT, S1, and S2 regions. Cases 1-4 represent different polydisperse aerosol configurations (see **Table 3** and **Fig. 4**), and their stagewise *DF* trends generally followed the corresponding stage cut-off diameters (*d*_50_) across the tested flow rates (see **Figs. 7 (a), 13 (a) and 13 (c)**). The discussion below focuses on MT, S1, and S2 because the preceding analysis showed that the most notable changes in combined stagewise *DF* occurred in this upstream region. Stages downstream of S2 were not emphasized because the influence of MT geometry became progressively weaker. For example, Case 1 showed less than 2% *DF* at S3 for all flow rates (see **Fig. 17(a)**), while Case 2 showed approximately 4-7% *DF* at S3, except for the USP IP case at 60 L/min, where the S3’s *DF* reached approximately 10% (see **Fig. 17(b)**). Similarly, for Cases 3 and 4, the deposition trends downstream of S3 showed only minor differences across flow rates and MT models (see **Figs. 16 (c) and (d)**). This behavior is consistent with the cascade nature of NGI deposition, i.e., particles not captured at an upstream stage would deposit at subsequent stages, causing the influence of MT geometry to diminish progressively downstream.

**Figure 17.**
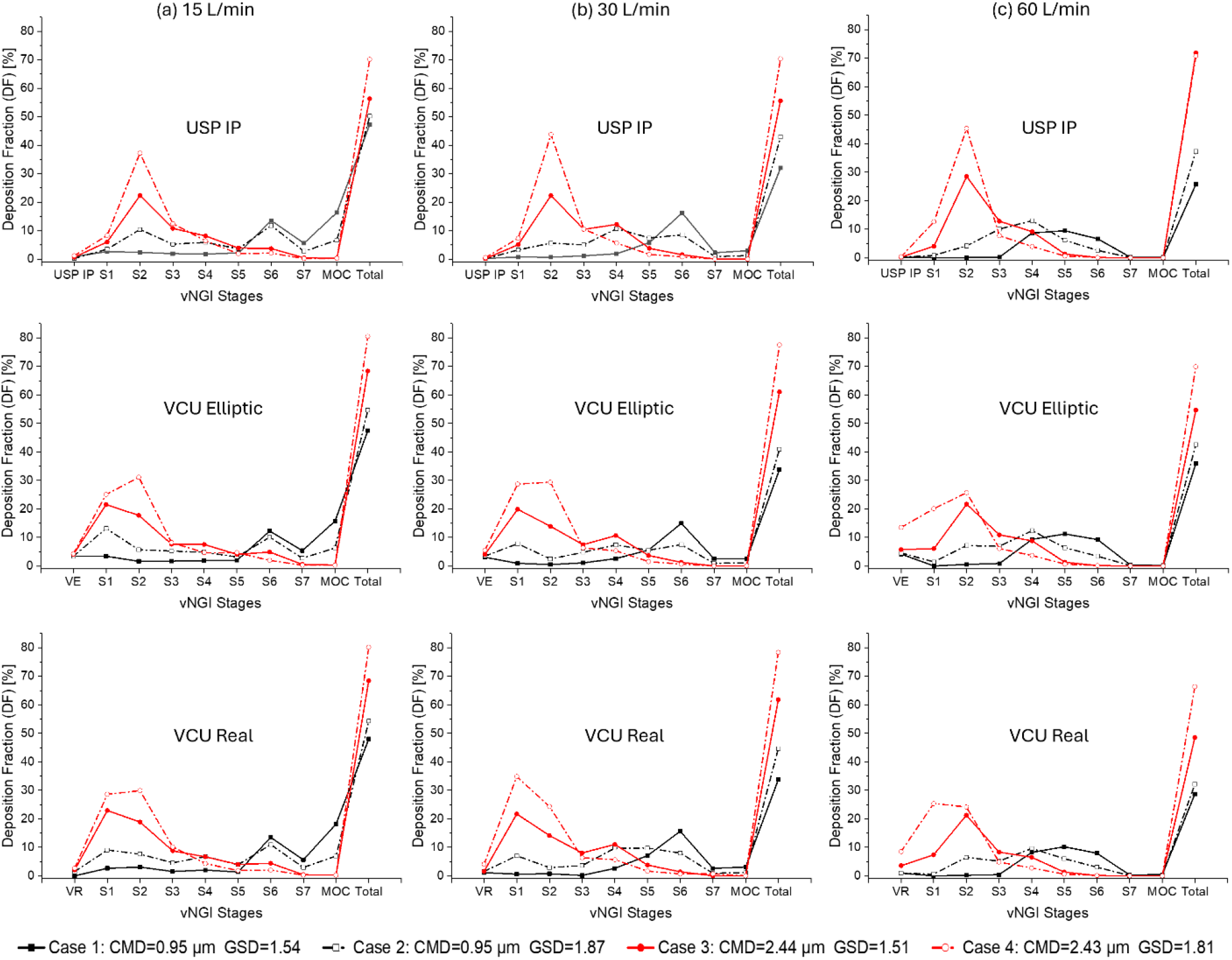
CFPD predicted regional deposition fractions (*DFs*) in the vNGI with three mouth-throat geometries, including the USP induction port (USP IP, top), VCU Elliptic (VE, middle), and VCU Real (VR, bottom), using four polydisperse particle size distributions (Cases 1-4) at three flow rates: (a) 15 L/min, (b) 30 L/min, and (c) 60 L/min.

When depositions in the MT region were examined separately, the USP IP collected approximately 1% or less of the particles and showed little sensitivity to flow rate. In contrast, the bio-relevant MT models showed clear dependence on both flow rate and aerosol size distribution (see **Table 6**). Both the VCU Real (VR) and VCU Elliptic (VE) MT models showed increasing *DF* with increasing flow rate. However, VR consistently produced lower MT deposition than VE with respective flow rates, consistent with previous observations under other aerosol conditions [42]. For Case 1 at 15 L/min, VR produced less MT *DF* comparable to that of the USP IP, whereas VE produced a MT *DF* approximately 3.8 times higher than that of the USP IP. This trend is consistent with experimental comparisons between the USP IP and VE geometries [75]. The effect became much stronger for larger particles and higher flow rates. For example, in Case 4 at 60 L/min, VR and VE collected 23.6 and 37.6 times as many particles as the USP IP, respectively, indicating a strong combined dependence on flow rate, particle size, and distribution breadth.

Both bio-relevant MT with Case 1, S1 and S2 each received less than 3.5% in *DF*, with higher deposition at 15 L/min and lower deposition at higher flow rates. This trend suggests that the smaller particles in Case 1 were able to follow the flow more effectively, with limited inertial impaction in the early stages. For Case 2, the *DF* in S2 generally decreased with increasing flow rate, although an exception was observed for the VR model at 30 L/min, where the *DF* in S2 dropped sharply to 2.9%. For the VE model in Case 2, the *DF* in S1 decreased with increasing flow rate, whereas the *DF* in S2 showed the opposite trend, except for the reduction observed at 30 L/min (see **Table 6**). Overall, the influence of bio-relevant MT geometry was less pronounced for Cases 1 and 2 than for Cases 3 and 4. The variations observed in Cases 1 and 2 may be associated with the combined effects of Brownian diffusion, local flow structures, and interstage wall loss, which requires further investigation in the future.

Cases 3 and 4 provide clearer evidence of the effect of MT geometry because these aerosols contained larger and heavier particles that were more susceptible to inertial impaction in the MT and early NGI stages (see **Fig. 4** and **Table 3**). This explains the higher *DF* observed in the bio-relevant MT models at increasing flow rates, with VE generally producing higher MT deposition than VR, and both bio-relevant MT models producing substantially higher deposition than the USP IP (see **Table 6**). In Case 3 both bio-relevant MT showed almost similar depositions at S1 and S2 at respective flow rates but the depositions in MT of VR are comparably less than VE. This trend could also be observed for total deposition for Case 3 and 4 with respective flow rates and more prominently for region wise in Case 3 than Case 4 (see **Table 5**). This indicates an approximate performance of VE with respect to VR in Case 3 and 4 much better than Case 1 and 2. In Case 4, deposition was distributed more evenly between S1 and S2 for the bio-relevant MT models (see **Table 6**), whereas deposition in the USP IP configuration remained concentrated around S2 (see **Fig. 16(d)**). Together, these observations demonstrate that bio-relevant MT geometry can significantly alter NGI performance for larger particles, particularly particles near or above approximately 4 µm, up to S2. The influence of MT geometry then decreased in the downstream stages, although it still affected the total *DF* to some extent (see **Table 5**). These findings highlight that changes in upstream inlet geometry and flow conditions can propagate downstream, modifying stagewise deposition.

The observed differences among MT models are likely to be related to geometry-induced changes in the airflow field and the resulting particle transport. Bio-relevant MT geometries contain anatomical features that can generate secondary flows and vortices within the MT [42]. As the flow rate increased, the turbulence intensity (*TI*) also increased, resulting in stronger turbulence dispersion and inertial impaction (see **Fig. 18**). Compared with the VE model, the VR model included multiple locations for jet impaction and regions of low *TI* where recirculation zones and eddies could develop. These features altered particle transport and deposition within the MT, leading to deposition behavior that differed substantially from that of the USP IP (see local deposition patterns in **Figs. 19-21**). As shown in **Figs. 19-21**, the MT configuration also affected the deposition pattern in S1. With the USP IP, particles tend to deposit preferentially on one side of S1, approximately opposite to the S1 outlet. In contrast, the bio-relevant MT models produced more spatially distributed deposition patterns. It can also be found that, at higher flow rates, S1 deposition became more concentrated near the MT jet-exit region because of stronger impaction on the S1 surface (see **Figs. 19-21**).

**Figure 18.**
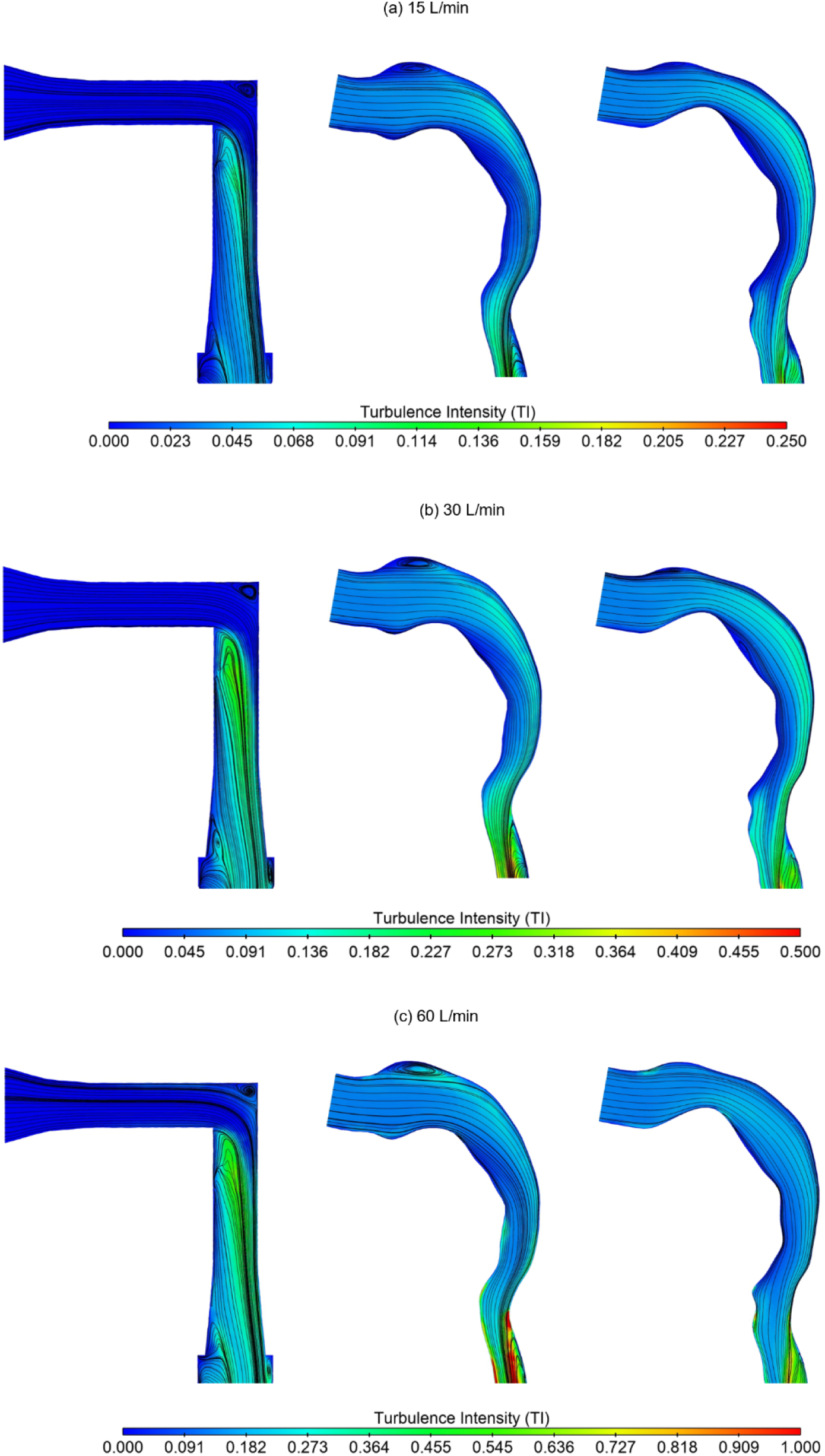
Turbulence intensity (*TI*) contour with velocity surface streamlines inside mid-plane of mouth-throat (MT) models, including USP IP (left), VCU Elliptic (middle), and VCU Realistic (right), at different flow rates: (a) 15 L/min, (b) 30 L/min, and (c) 60 L/min

**Figure 19.**
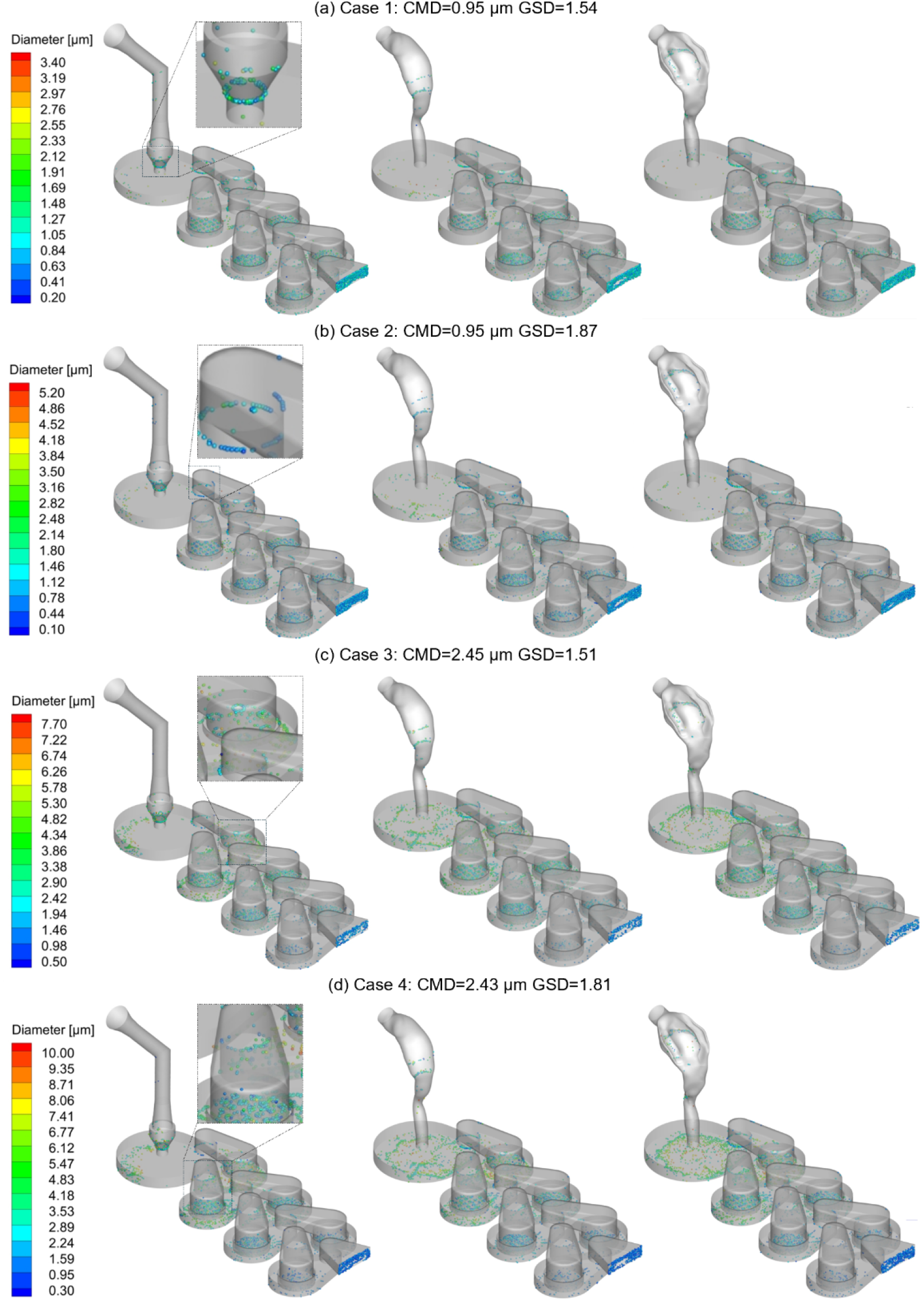
Localized polydisperse particle deposition patterns with insets of common interstage wall losses (*WL*) areas in vNGI with different mouth-throat geometries (USP IP (left), VCU Elliptic (middle), VCU Real (right)) at 30 L/min: (a) Case 1, (b) Case 2, (c) Case 3, and (d) Case 4

**Figure 20.**
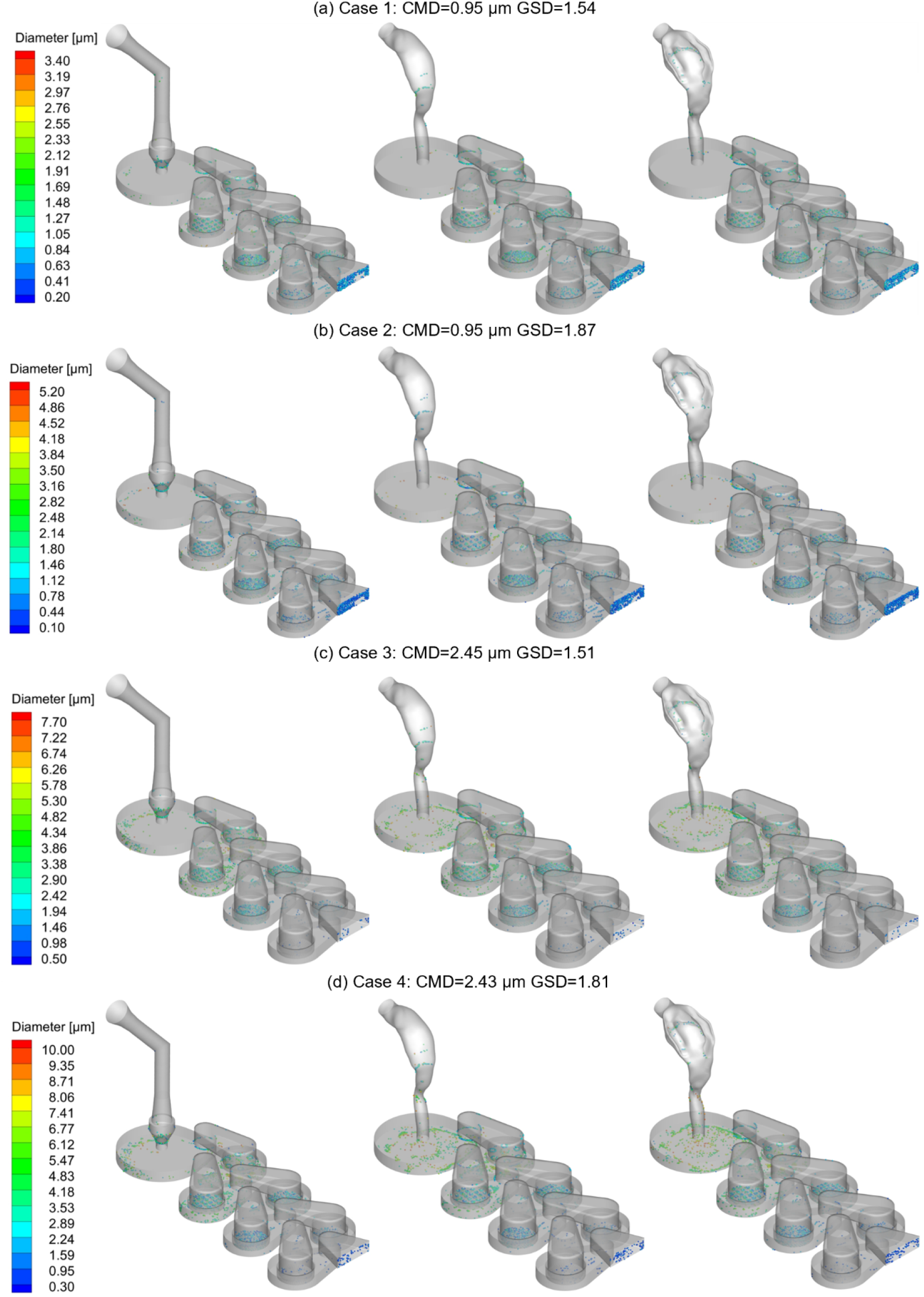
Localized polydisperse particle deposition patterns in vNGI with different mouth-throat geometries (USP IP (left), VCU Elliptic (middle), VCU Real (right) at 30 L/min: (a) Case 1, (b) Case 2, (c) Case 3, and (d) Case 4

**Figure 21.**
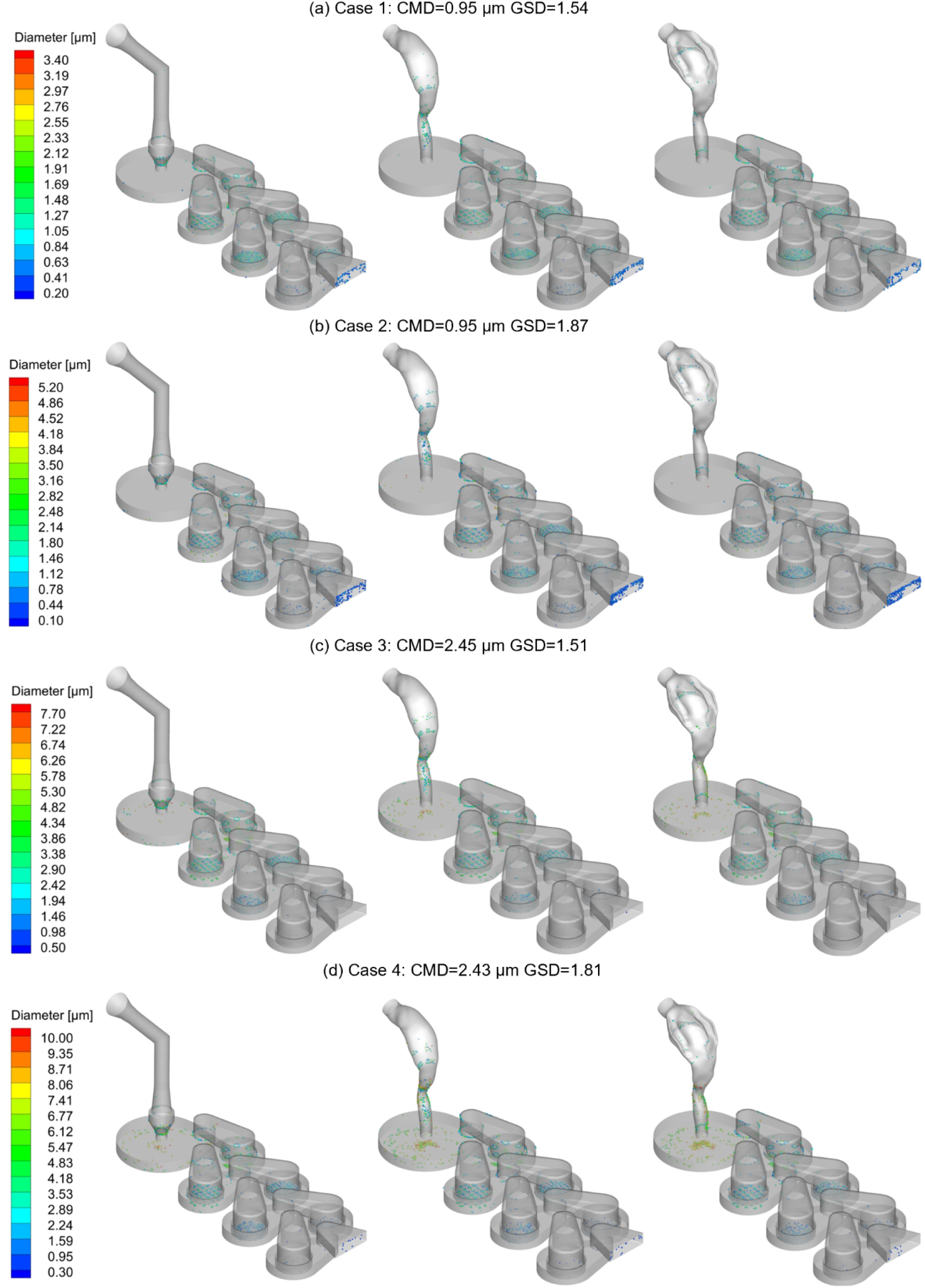
Localized polydisperse particle deposition patterns in vNGI with different mouth-throat geometries (USP IP (left), VCU Elliptic (middle), VCU Real (right) at 60 L/min: (a) Case 1, (b) Case 2, (c) Case 3, and (d) Case 4

The difference in local deposition patterns induced by the choice of MT configuration is due to the differences in the airflow structures exiting the MT and entering S1. In the USP IP case (see **Fig. 18**), a large circulation region was observed near the lower left corner, whereas in the bio-relevant MT models, anatomical features such as the epiglottis generated smaller eddies near the lower-right side of the outlet region, consistent with previous studies [42]. These differences in MT exit-flow patterns, together with the flow development inside S1 before the stage outlet, produced the observed differences in S1 deposition patterns and the increased *DF* in S1 with the bio-relevant MT models compared with the USP IP (see **Table 6**). Overall, these coupled air-particle interactions altered the effective cutoff behavior of the MT and S1 regions, while their influence progressively diminished in the downstream vNGI stages.

## 4. Conclusions

This study developed a computational fluid particle dynamics (CFPD)-based virtual Next Generation Impactor (vNGI) to resolve airflow structures and particle transport/deposition behavior within the NGI under various flow rates, particle size distributions, and MT configurations. The vNGI enables spatiotemporal analysis of particle-laden flows in regions that are difficult to quantify experimentally. In addition to predicting stagewise deposition behavior, the model provides mechanistic insight into crossflow, interstage *WL*, flow-rate-dependent deposition, and the influence of bio-relevant MT geometries. These capabilities support the potential use of the vNGI as a complementary *in silico* NAM for inhalation product development and APSD measurement. The major conclusions are summarized as follows.

1. A CFPD-based vNGI provides a mechanistically informative digital representation of the NGI for APSD-related analysis. Because APSD is a critical quality attribute for orally inhaled drug products, conventional NGI experiments often require repeated testing under product-specific flow rates and environmental conditions. The vNGI developed in this study can complement *in vitro* experiments by enabling controlled, repeatable, and high-resolution analysis of airflow and aerosol deposition behavior under different operating conditions.
2. The vNGI was reasonably validated against available experimental data at a steady flow rate of 30 L/min using stage cut-off diameter, *d*_50_, and collection efficiency behavior. The model captured the general stagewise deposition trend, with stronger predictive performance in the earlier and intermediate NGI stages. Discrepancies in the downstream stages, particularly S6 and S7, were associated with known limitations of the point-particle DPM approach, the omission of the fully resolved MOC geometry, and uncertainty in wall-deposition/re-entrainment behavior. Therefore, the current vNGI should be interpreted as a fit-for-purpose mechanistic tool rather than a direct replacement for all experimental NGI measurements.
3. The airflow analysis showed that crossflow remains an important design and performance consideration in multi-nozzle inertial impactor stages. Crossflow was more pronounced near the central region of the nozzle filter than near the peripheral region. The results suggest that conventional geometric criteria, such as the ratio of jet-to-plate distance to nozzle diameter, may not be sufficient to fully characterize or eliminate crossflow in complex multi-nozzle stages. A progressive nozzle-pitch arrangement, while maintaining the same overall nozzle-cluster diameter, may help reduce crossflow effects and improve impactor-stage performance.
4. Interstage *WL* was strongly influenced by particle size. For monodisperse aerosols, particles smaller than approximately 1 µm contributed substantially to interstage *WL*, thereby reducing the total stagewise deposition fraction. These findings indicate that submicron particles are more susceptible to deposition outside the intended collection stages, likely due to the combined effects of Brownian-motion-induced diffusion, turbulent dispersion, and local flow separation in interstage regions.
5. Flow rate affected stagewise deposition differently depending on the aerosol size distribution. For smaller polydisperse aerosols with *CMD* ≈ 0.95 μ*m* The stagewise deposition fraction was more sensitive to changes in flow rate, with deposition shifting among downstream stages as the stage cut-off diameters changed. In contrast, larger polydisperse aerosols with *CMD* ≈ 2.4 − 2.5 μ*m* were less sensitive to flow-rate variation in total deposition, although their stagewise deposition shifted toward upstream NGI stages due to increased inertial impaction.
6. Bio-relevant MT geometries altered stagewise deposition patterns, particularly in the MT, S1, and S2 regions. Compared with the USP IP, the VCU Elliptic and VCU Realistic MT models exhibited distinct upstream deposition behavior due to geometry-induced secondary flows, vortices, and altered jet-exit conditions at the entry to S1. These effects were most evident for larger aerosol distributions, where particles were more susceptible to inertial impaction in the MT and early NGI stages. Although the total *DF* differences among MT models were sometimes modest, the stagewise redistribution caused by MT geometry was significant and should be considered when interpreting APSD measurements under more bio-relevant inlet conditions.
7. The influence of MT geometry decreased progressively in the downstream NGI stages. This behavior is consistent with the cascade nature of the NGI: particles not captured in an upstream region would deposit in subsequent stages, causing the upstream geometric effect to diminish downstream. Nevertheless, changes in MT geometry and upstream flow structure can propagate into S1 and S2, thereby modifying the effective stagewise deposition behavior.

Overall, vNGI provides a flexible and extensible *in silico* platform for evaluating how flow rate, aerosol size distribution, interstage *WL*, and MT geometry influence APSD-related deposition behavior. With further refinement and expanded validation, including improved treatment of particle-wall interactions, re-entrainment, finite-size particle effects, and MOC representation, the vNGI could support inhalation product development, device optimization, formulation screening, and regulatory-relevant alternative bioequivalence assessment as a complementary NAM alongside conventional *in vitro* NGI testing.

## 5. Limitations of This Study and Future Work

Despite the capability of predicting air-particle flow dynamics using the CFPD-based vNGI developed in this study and the insights generated from the high-resolution spatiotemporal distributions of variables of interest, a couple of limitations remain, i.e.,

1. The DPM model treats particles as point masses and therefore does not explicitly resolve finite-size effects. This limitation may reduce prediction accuracy when particle diameters are comparable to the nozzle/filter dimensions, especially in the downstream NGI stages. Additionally, particle-particle interactions were also neglected under the dilute-aerosol assumption; however, under higher particle-loading conditions, collisions, agglomeration, and momentum exchange may influence particle transport, stage-to-stage penetration, and deposition behavior.
2. The last MOC stage of NGI was not considered for CFPD modeling, which may influence the S6 and S7 fluid flow, and in turn the *CE* or *DF*.
3. Appropriate boundary conditions for particle depositions, such as partial trap boundary conditions to mimic bounce and re-entrainment of particles based on an appropriate coefficient of restitution at walls other than the collection stages, were not implemented due to a lack of available literature on interstage wall losses.
4. A limited number of particles were considered for polydisperse aerosols.

To address the above-mentioned limitations, future work may include:

1. Incorporation of finite-size particle modeling approaches (e.g., resolved particle methods or corrections to point-particle assumptions) to improve the accuracy of particle transport predictions, particularly for submicron and transitional diameter ranges where continuum assumptions may break down. Integration of particle– particle interaction models (e.g., discrete element method (DEM) or stochastic collision models) to capture agglomeration, collision, and transition behaviors of larger aerosol particles under high particle loading conditions.
2. Implementation of mesh refinement strategies coupled with improved particle–fluid coupling schemes, enabling grid-independent solutions while mitigating numerical artifacts associated with the current DPM framework.
3. Extension of the computational domain to include the final MOC stage of the NGI, allowing for more accurate representation of downstream flow structures and their influence on particle collection efficiency (*CE*) or deposition fraction (*DF*) in S6 and S7.
4. Development and validation of advanced wall interaction and deposition boundary conditions, particularly for interstage regions, through a combination of in silico studies and targeted experimental measurements to better quantify and model interstage wall losses.
5. Consideration of an adequate number of particles per particle diameter size bins for polydisperse aerosol based on the particle number independence test or the realistic particle number injected based on real-world particle mass flow rate.

## Nomenclature

**Acronyms**

ACI: Anderson Cascade Impactor
APRD: Average Percentage of Relative Difference
APSD: Aerodynamic Particle Size Distribution
CAD: Computer-Aided Design
CE: Collection Efficiency
CFD: Computational Fluid Dynamics
CFPD: Computational Fluid Particle Dynamics
CMD: Count Median Diameter
DF: Deposition Fraction
DPM: Discrete Phase Model
GEKO: Generalized k - ω
GSD: Geometric Standard Deviation
IP: Induction Port
MMAD: Mass Median Aerodynamic Diameter
MMD: Mass Median Diameter
MOC: Micro-Orifice Collector
MT: Mouth-throat
NAMS: New Approach Methodologies
NGI: Next Generation Impactor
OIDPs: Orally Inhaled Drug Products
PSG: Product Specific Guidelines
RANS: Reynolds Averaged Navier-Stokes
SAC: Single Actuation Content
SIMPLE: Semi-implicit pressure linked equation
SMI: Soft Mist Inhaler
TI: Turbulence Intensity
USP: U.S. Pharmacopeia
VE: VCU Elliptic
vNGI: Virtual Next Generation Impactor
VR: VCU Realistic
WL: Wall Loss
WSS: Wall Shear Stress

**Symbols**

*λ*: Air Mean Free Path
*C*_*c*_: Cunningham Slip Correction Factor
*D*_*c*_: Diameter of the Cluster of Nozzles on a Stage
*d*_*p*_: Particle Diameter
*d*_*ae*_: Aerodynamic Particle Diameter
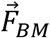: Brownian Motion-Induced Force
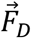: Drag Force
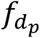: Particle Number Distribution
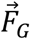: Gravitational Force
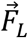: Lift Force
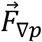: Pressure Gradient Force
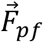: Aerodynamic Forces Acting on Particles
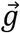: Gravitational Acceleration
*h*: Cup Height of a Stage
*k*: Turbulent Kinetic Energy
*K*_*n*_: Knudsen Number
*m*_*p*_: Particle Mass
*n*: Number of Nozzles in a Stage
*N*_*down*_: Number of Particles Deposited Downstream
*N*_*Total*_: Total number of injected particles
*N*_*up*_: Number of Particles Deposited Upstream
*p*: Pressure
*P*: Pitch (Distance or Angle between Two Nozzles)
*Q*: Volumetric Flow Rate
*Re*: Reynolds Number
*S*: Jet-to-plate Distance
*S*_*t*_: Stokes Number
*t*: Time
*u*: Fluid Phase Velocity Magnitude
*u*_*p*_: Particle Velocity
∀: Volume
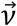: Velocity Vector
*v*_*o*_: Velocity Magnitude at the Nozzle Exit
*ω*: Specific Dissipation Rate
*W*: Filter Nozzle Diameter in the vNGI stages
*μ*: Viscosity
*ρ*: Fluid Phase Density
*ρ*_*ae*_: Aerodynamic Particle Density
*ρ*_*p*_: Particle Density
*σ*_*κ*_, *σ*_*ω*_: Prandtl numbers for *k* and *ω*
*τ*_*r*_: Particle Relaxation Time
*X*_*c*_: Cross-flow Parameter

## Acknowledgment

This work acknowledges Dr. Lucila Garcia-Contreras from the University of Oklahoma Health Science Center, Oklahoma City, OK, USA, for facilitating the measurement of NGI and providing technical guidance. The authors also thank Dr. Hang Yi in Chemical Engineering at Oklahoma State University for his assistance with figure preparation. This work was supported by the U.S. National Science Foundation (NSF) under Award No. TI-2234619. The use of Ansys software (Ansys Inc., Canonsburg, PA, USA) as part of the Ansys-CBBL academic partnership agreement is gratefully acknowledged (Thierry Marchal and Vishal Ganore). The simulations were performed in the PETE supercomputer funded by MRI Award #1531128, Acquisition of Shared High Performance Compute Cluster for Multidisciplinary Computational and Data-Intensive Research.

## Data availability

The data that support the findings of this study are available from the corresponding author upon reasonable request.

